# Structural analysis of Red1 as a conserved scaffold of the RNA-targeting MTREC/PAXT complex

**DOI:** 10.1101/2022.01.19.476877

**Authors:** Anne-Emmanuelle Foucher, Leila Touat-Todeschini, Ariadna B. Juarez-Martinez, Hamida Laroussi, Claire Karczewski, Samira Acajjaoui, Montserrat Soler-López, Stephen Cusack, Cameron D. Mackereth, André Verdel, Jan Kadlec

**Affiliations:** Univ. Grenoble Alpes, CNRS, CEA, IBS, F-38000 Grenoble, France; Institut for Advanced Biosciences, UMR Inserm U1209/CNRS 5309/University Grenoble Alpes, La Tronche, France; Structural Biology Group, European Synchrotron Radiation Facility (ESRF), CS 40220, 38043 Grenoble, France; European Molecular Biology Laboratory, Grenoble Outstation, 71 Avenue des Martyrs, CS 90181, Grenoble Cedex 9 38042, France; Univ. Bordeaux, Inserm U1212, CNRS UMR 5320, ARNA Laboratory, Institut Européen de Chimie et Biologie, 33607 Pessac, France

## Abstract

To eliminate specific or aberrant transcripts, eukaryotic cells use nuclear RNA-targeting complexes that deliver them to the exosome for degradation. *S. pombe* MTREC complex, and its human counterpart PAXT, are key players in this mechanism. Red1 and hZFC3H1 function as scaffolds of these respective complexes. Here, we present an NMR structure of a helix-turn-helix domain of Red1 in complex with the N-terminus of Iss10 and show this interaction is required for proper cellular growth and meiotic mRNA degradation. We also report a crystal structure of a Red1*-*Ars2 complex that explains the mutually exclusive interactions of hARS2 with various “ED/EGEI/L” motif-possessing RNA regulators such as hZFC3H1, hFLASH or hNCBP3. Finally, we show that both Red1 and hZFC3H1 homo-dimerize via their coiled-coil regions indicating that MTREC and PAXT likely function as dimers. Our results, combining structures of three Red1 binding interfaces with *in vivo* studies, provide mechanistic insights into conserved features of MTREC/PAXT architecture.

## Introduction

An astonishing amount of RNA, including a sizeable portion of aberrant transcripts, is constantly being synthesised within eukaryotic nuclei^1–3^. To detect and eliminate specific or aberrant transcripts, cells rely on highly regulated RNA degradation machineries. Unwanted transcripts are recognised by specialised RNA-targeting complexes that deliver them to the nuclear RNA exosome for degradation^4, 5^.

In the fission yeast *Schizosaccharomyces pombe*, MTREC (Mtl1–Red1 core) or NURS (nuclear RNA silencing) is such an RNA-targeting complex and is responsible for recognition of cryptic unstable transcripts (CUTs), meiotic mRNAs and unspliced pre-mRNAs^6–8^. The MTREC core module comprises the zinc-finger protein Red1 and the RNA helicase Mtl1. In addition, there are four associated sub-modules - the nuclear cap-binding complex (Cbc-Ars2), the Iss10-Mmi1 complex that targets meiotic RNAs, the Pab2-Rmn1-Red5 complex likely recognizing poly(A) sequences and the more loosely associated poly(A) polymerase Pla1^6–8^. The subunit conservation indicates that MTREC is the fission yeast counterpart to the human PAXT (pA-tail-exosome-targeting) complex that targets long and polyadenylated nuclear RNAs^7, 9–11^. Indeed, PAXT contains a large zinc-finger protein ZFC3H1 and the RNA helicase MTR4, human homologs of MTREC Red1 and Mtl1, as well as the cap-binding and poly(A)-binding sub-modules^9–11^. The well-studied human NEXT (nuclear exosome-targeting) complex, on the other hand, is composed of the MTR4 helicase, zinc-knuckle protein ZCCHC8 and the RNA-binding subunit RBM7^1^^2^. NEXT is involved in degradation of short-lived RNAs, such as promoter upstream transcripts (PROMPTs) and enhancer RNAs (eRNAs)^12, 13^ but its relation to MTREC is less clear.

The subunit composition of the *S. pombe* MTREC and human PAXT complexes has been characterized by proteomics^6–9, 11^. In MTREC, the Red1 protein was suggested to form a scaffold of the complex, interacting with multiple subunits and linking individual sub-modules together^6, 7, 14^. Red1 consists of 711 residues (compared to the 1989 residues of its ZFC3H1 counterpart in human PAXT) and is predicted to be mostly disordered. Red1 interacts with the Mtl1 Arch domain via its zinc-finger containing domain at its C-terminus^14^. In human and *S. cerevisiae*, Mtl1 counterpart MTR4 has been structurally characterised^15^ revealing molecular details of its interactions with its partners including the NEXT subunit ZCCHC8^16^ and the RNA exosome to whom MTR4 presents the targeted RNA for degradation^17, 18^. How the Red1-Mtl1 core interacts with the individual MTREC sub-modules and what is their function in MTREC/PAXT activity, however, remains unclear.

The MTREC complex is involved in specific elimination of meiotic transcripts and heterochromatin formation in *S. pombe* and this activity involves the Iss10/Mmi1 sub-module^6–8, 19, 20^. Indeed, in *S. pombe*, expression of a number of meiotic genes is strictly repressed during the vegetative growth by selective RNA degradation, since their aberrant expression is highly deleterious to cell viability^21^. Meiotic transcripts and non-coding RNAs that contain repeats of a hexanucleotide motif called determinant of selective removal (DSR) are specifically recognized by the YTH family RNA-binding protein Mmi1, which in turn recruits MTREC to mediate their degradation^20–24^. In vegetatively growing cells, Mmi1 forms nuclear foci^21^, where most of its partners, including MTREC, nuclear exosome and the target RNA, co-localize^6, 8, 25^. The Iss10 subunit is also directly involved in the selective elimination of meiotic DSR-containing transcripts and is important for the MTREC sub-nuclear localization^26^. In addition, Iss10 and the MTREC-dependent RNA degradation has been connected to the mTOR cellular energy-sensing signalling^27^. Under vegetative growth, Iss10 is phosphorylated by TORC1 (TOR complex 1), which prevents its degradation by the proteasome. However, under conditions of nitrogen depletion, TORC1 phosphorylation of Iss10 is blocked and Iss10 in turn eliminated^27^. Lack of Iss10 comes with a loss of MTREC co-localization with Mmi1 in nuclear foci, and accumulation of DSR-containing mRNAs and meiotic proteins, thereby allowing the cells to proceed with meiosis^26, 27^. Thus, Iss10 is a key regulatory element of MTREC-dependent degradation of meiotic mRNAs, that most likely links the MTREC core complex to the Mmi1 bound RNAs^26^.

The Red1-Mtl1 core also recruits the Cbc-Ars2 sub-module^6, 7^, which has been more extensively studied in human. CBC (nuclear cap-binding complex) is a key RNA biogenesis regulator that co-transcriptionally binds the m^7^G cap at the 5’ end of RNA polymerase II transcripts^28, 29^. The CBC and its partner ARS2 are implicated in fate determination of Pol II transcripts, including their processing, transport or degradation^30^. CBC-ARS2 interacts in a mutually exclusive way with positive RNA biogenesis factors (such as PHAX, FLASH and NCBP3), or with the RNA destructive MTREC/PAXT and NEXT assemblies^30–32^. While the connection to PAXT and NEXT involves ZC3H18^9^, no such protein has been identified in the MTREC complex^6, 7^. How the CBC-ARS2 complex is integrated and what role it plays in the exosome-linked RNA targeting complexes is currently unknown.

Here, we report a biochemical and structural analysis of two important interactions of Red1 with MTREC sub-modules, combined with *in vivo* studies. We mapped minimal Red1 regions required for its interaction with Iss10 and Ars2, respectively. We determined an NMR structure of the Red1-Iss10 complex, which reveals how Red1 recognises Iss10 via its helix-turn-helix motif. We also report a crystal structure of the Red1-Ars2 complex, which shows how the Ars2 C-terminal zinc-finger domain interacts with the conserved EDGEI motif of Red1. In addition, we demonstrated equivalent interaction between ZFC3H1 and hARS2 within the PAXT complex, providing the first structural insights into how the Cbc-Ars2 complex is integrated into the RNA-targeting complexes. Functional *in vivo* experiments using Red1 structure-driven mutants showed that both Iss10 and Ars2 interactions are important for MTREC mediated cellular activities. Finally, we demonstrated that Red1 dimerizes via its coiled-coil region, strongly suggesting that the MTREC complex acts as a dimeric assembly and this feature is conserved in human PAXT and NEXT complexes.

## Results

### Red1 forms multiple interactions within the MTREC complex

Red1, together with the Mtl1 helicase, has been proposed to form the MTREC core, but the molecular details of its interactions with the MTREC sub-modules remain unknown (Fig. 1a). To understand how Red1 associates with its partners, we first performed yeast two-hybrid (Y2H) assays to analyse its interactions with several MTREC members. Using Red1 as bait, the expected interaction with Mtl1 could be detected (Fig. 1b). In addition, Red1 also strongly interacted with Iss10 and Ars2 (Fig. 1b). Both the Ars2 and Iss10 interactions were confirmed in an inverted experiment where Red1 was used as a prey protein (Fig. 1c,d). These assays also showed a complex formation between Iss10 and Mmi1 (Fig. 1c). Finally, the Y2H assay revealed an unexpected Red1 self-association (Fig. 1e). These results thus indicate that Red1 forms at least three distinct and direct interactions with other MTREC subunits and has the potential to oligomerize (Fig. 1f). This suggests that Red1 acts as the MTREC scaffold protein.

**Figure 1.**
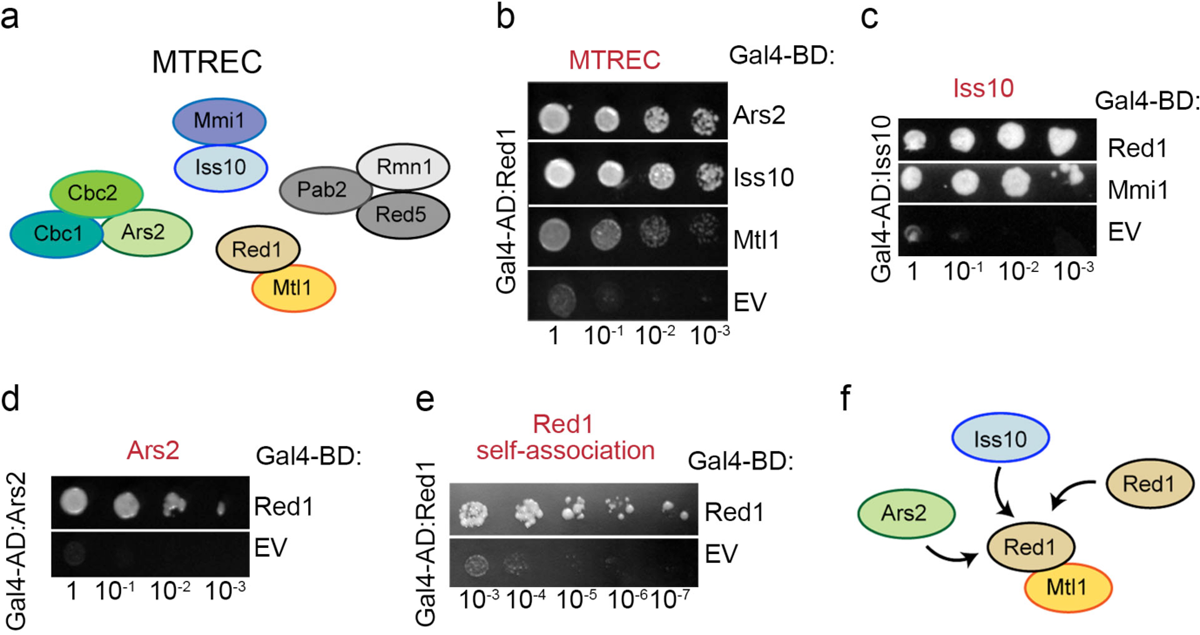
Red1 forms multiple interactions within the MTREC complex. **(a)** Schematic representation of the MTREC complex. Red1 and Mtl1 form the MTREC complex core^7^. **(b)** Y2H assays revealed that Red1 interacts with Ars2 and Iss10. Mtl1 was used a positive control. Red1 was used as bait. Growth assayed on selective medium lacking histidine and with 50mM 3-amino-1,2,4-triazole (3AT) histidine inhibitor is shown. EV-empty vector. **(c**) Y2H assays with Red1 as a prey showing that Iss10 interacts with Red1. Iss10 interacts also with Mmi1. Growth was assayed on medium lacking histidine and with 20mM and 30mM 3AT. **(d)** Y2H assays with Red1 as a prey showing that Ars2 interacts with Red1. Growth was assayed on medium lacking histidine and with 20mM 3AT. **(e)** Y2H analysis of Red1 self-association. Growth was assayed on medium lacking histidine and with 30mM 3AT. **(f)** Schematic representation of the interactions characterised in this study.

### Red1 interacts with the N-terminus of Iss10

In order to define the interacting regions between Red1 and Iss10, both proteins were split into three consecutive fragments and their interactions tested by Y2H. While a clear interaction was observed for the N-terminal part of Iss10 (1-152) and the middle part of Red1 (138-390), the remaining constructs did not interact (Supplementary Fig. 1a,b). The binding of these two fragments was confirmed *in vitro* by Strep-tag pull-down experiments where Red1^138–390^ could be co-purified with Iss10^1–152^, requiring co-expression of the two proteins (Supplementary Fig. 1c, lane 4). Several deletions based on limited proteolysis and sequence alignments were produced, eventually yielding an Iss10^1-45^ construct that retained its capacity to bind Red1^138-390^ (Supplementary Fig. 1c, lane 5). Red1 could also be reduced to the minimal Iss10-binding fragment spanning residues 192-236 (Supplementary Fig. 1c, lane 6). Furthermore, Multi angle laser light scattering (MALLS) measurements revealed a 1:1 hetero-dimer (Supplementary Fig. 1d). In summary, these results show that Red1 and Iss10 interact together via short regions within Red1^192-236^ and Iss10^1-45^ (Fig. 2a).

**Figure 2.**
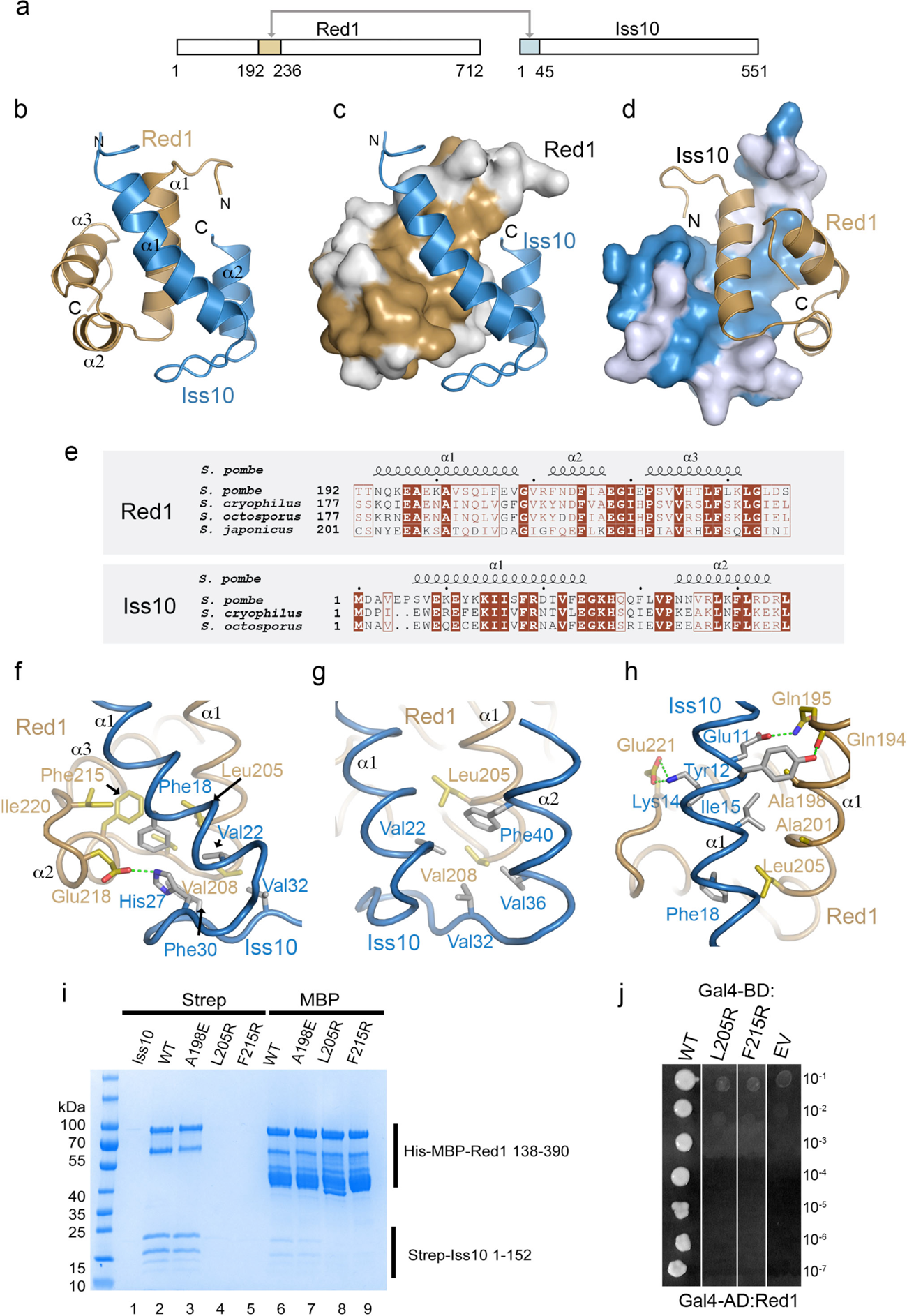
Red1 interacts with Iss10 with its HTH domain. **(a)** A schematic diagram highlighting the Red1 and Iss10 interacting regions as defined in this study. **(b)** Ribbon representation of the lowest energy NMR-based structure of the Red1-Iss10 complex. The Red1 HTH domain is shown in brown and the Iss10 N-terminus is in blue. The secondary structures are labeled. **(c)** Surface representation of Red1 with conserved surface residues shown in brown. **(d)** Surface representation of Iss10 with conserved surface residues highlighted in blue. **(e)** Sequence alignment of the Red1 HTH domain (upper panel) and Iss10 (lower panel) in different *Schizosaccharomyces* species. Identical residues are in brown boxes. **(f)** Details of the interaction between the Iss10 helix α1 and the following loop with Red1 helices α1 and α3. **(g)** Details of the Red1 helix α1 packing against the Iss10 surface made of helices α1 and α2. **(h)** Coiled-coil interaction and hydrogen bonding between Red1 and Iss10. **(i)** Strep-tag pull-down experiments with Strep-Iss10^1-45^ and His-MBP-Red1^192-236^ mutants indicated above the lanes. Iss10^1-45^ and Red1^192-236^ were co-expressed, the cultures were split into two halves and the proteins purified on Strep-tactin or amylose resins. In absence of the interaction with Red1, Strep-Iss10^1-45^ is unstable. **(j)** Essential role of Red1 Lue205 and Phe215 in the interaction with Iss10 shown by Y2H assay. Growth was assayed on medium lacking histidine and with 30mM 3AT.

### NMR structure of the Red1-Iss10 complex

The Red1^192-236^-Iss10^1-45^ complex was then subject to structural analysis. While the complex resisted crystallization, we obtained a high quality ^15^N-HSQC NMR spectrum (Supplementary Fig. 1e) that prompted us to determine its structure by NMR. The resulting structural ensemble (Supplementary Table 1; Supplementary Fig. 2a,b) revealed that Red1 residues 192-236 and Iss10 residues 1-44 comprise the structured core of the interaction complex. Red1^192-236^ folds into three short helices arranged in a way reminiscent to the classical helix-turn-helix (HTH) motif (Fig. 2b-d, Supplementary Fig. 2c). Indeed, a search of the Protein Data Bank using the PDBeFold server at the EBI revealed that Red1^192-236^ is highly similar to other HTH motifs, such as the DNA binding domain of the *B. subtilis* DeoR protein^33^ (Supplementary Fig. 2c-e). Nevertheless, the surface of Red1 helix α3, that in classical HTH motifs frequently mediates dsDNA binding, carries mostly hydrophobic residues (Supplementary Fig. 2f). While Red1 DNA binding has not been reported and no unspecific DNA binding was observed during Red1 purification, DNA interaction cannot be excluded. The Iss10^1-45^ fragment consists of two helices, packing tightly against Red1 (Fig. 2b-d). Mutual interaction surfaces on both proteins are made of residues that are highly conserved among evolutionary distant *Schizosaccharomyces* species underlining the importance of this interaction (Fig. 2e).

### The Red1-Iss10 interface

The interface between Red1 and Iss10 is dominated by hydrophobic interactions. Iss10 helix α1 and the following loop form numerous contacts with the groove between Red1 helices α1 and α3 (Fig. 2f). Iss10 Phe18, Val22, Phe30, and Val32 interact with the Red1 hydrophobic surface composed of Leu205, Val208, Val210, Phe215 and Ile220. Glu218 also makes a salt bridge with His27 (Fig. 2f). In parallel, Red1 helix α1 packs against the Iss10 surface made of helices α1 and α2. Red1 Leu205 and Val208 interact with Iss10 residues around Val22, Val32, Val36 and Phe40 (Fig 2g). Finally, Iss10 and Red1 establish a short coiled-coil interaction, including contacts between Iss10 Ala198, Ala201 and Leu205 and Red1 Ile15 and Phe18 (Fig. 2h). These interactions are further strengthened by several hydrogen bonds of Iss10 Glu11, Tyr12 and Lys14 and Red1 Gln194, Gln195 and Glu221 (Fig. 2h).

We were next interested in identifying Red1 mutations that could disrupt its interaction with Iss10 without affecting the Red1 HTH fold. To this end, we mutated Ala198 in the centre of the coiled-coil formed with Iss10 and two hydrophobic residues Leu205 and Phe215 that we hypothesised should directly disrupt the interaction with Iss10 helix α1 (Fig. 2f-h). These mutations were introduced into the His-MBP-Red1^138-390^ and tested for their interaction with Iss10^1-152^ by using Strep- and MBP-pull-down assays (Fig. 2i). While A198E retained the WT level of binding (Fig. 2i, lanes 3 and 7), the L205R and F215R mutants no longer bound Iss10 (Fig 2i, lanes 4,5,8 and 9). Since co-expression with Red1 is required for Iss10^1-152^ stability, Iss10^1-152^ produced with these Red1 mutants becomes unstable and does not purify on the Strep-Tactin resin. We then confirmed the essential role of Leu205 and F215R for the interaction with Iss10 in the context of full-length proteins using Y2H assay (Fig. 2j). Ultimately, the L205R mutation was selected for *in vivo* studies (see below) and the integrity of mutated Red1^192-236^ was verified by gel filtration, where its elution profile was similar to the WT protein (data not shown).

### Red1-Iss10 binding interface is required for proper degradation of meiotic DSR-containing mRNAs and clustering of Red1 foci *in vivo*

In order to evaluate *in vivo* the function of Red1-Iss10 binding interface, recombinant Red1-TAP or Red1-L205R-TAP *proteins* in combination with Iss10-GFP protein were expressed in *S. pombe cells* in place of the endogenous proteins using the respective endogenous promoters. Using these cells, co-immunoprecipitation experiments showed that the Red1-L205R mutation compromised Red1 association with Iss10 *in vivo* (Fig. 3a). Next, we examined the growth of *red1-L205R* mutant cells. While *red1Δ* cells showed only a moderate growth defect on solid rich medium at 30°C, this defect was more pronounced at lower temperature of 25 or 18°C (Supplementary Fig. 3a) or when the cells were grown on minimal medium (Supplementary Fig. 3b). Importantly, *red1-L205R* cells showed a similar growth defect, when grown on minimal medium, suggesting that Red1-Iss10 interaction can be important for Red1-dependent cell growth function depending on the environmental conditions (Fig. 3b). We then examined the subcellular localization of Red1-L205R-TAP mutant and Iss10-GFP. Both Red1 and Iss10 have been previously reported to co-localize within nuclear foci^23,26,34^. We observed that the *red1-L205R* mutation had an effect on both Iss10 and Red1 localization by, respectively, inducing the disappearance of Iss10 nuclear foci and the increase in the number of Red1 foci per nucleus (Fig. 3c-e). These findings indicate that the interaction of Iss10 with Red1 is required for its recruitment to Red1 nuclear foci and suggest that the Iss10 and Red1 interaction may promote the clustering of Red1 nuclear foci. Finally, we examined the level of meiotic DSR-containing mRNAs which are degraded by the MTREC machinery^7^. Expression of the Red1-L205R mutant, instead of WT Red1, led to a defect comparable to *iss10Δ* mutation and it was more pronounced when cells were grown in minimal medium (Fig. 3f and Supplementary Fig. 3c). Taken together, we conclude that *in vivo*, Red1-Iss10 binding interface plays a role in mediating functions of Red1, and its importance is determined by the environmental growth conditions.

**Figure 3.**
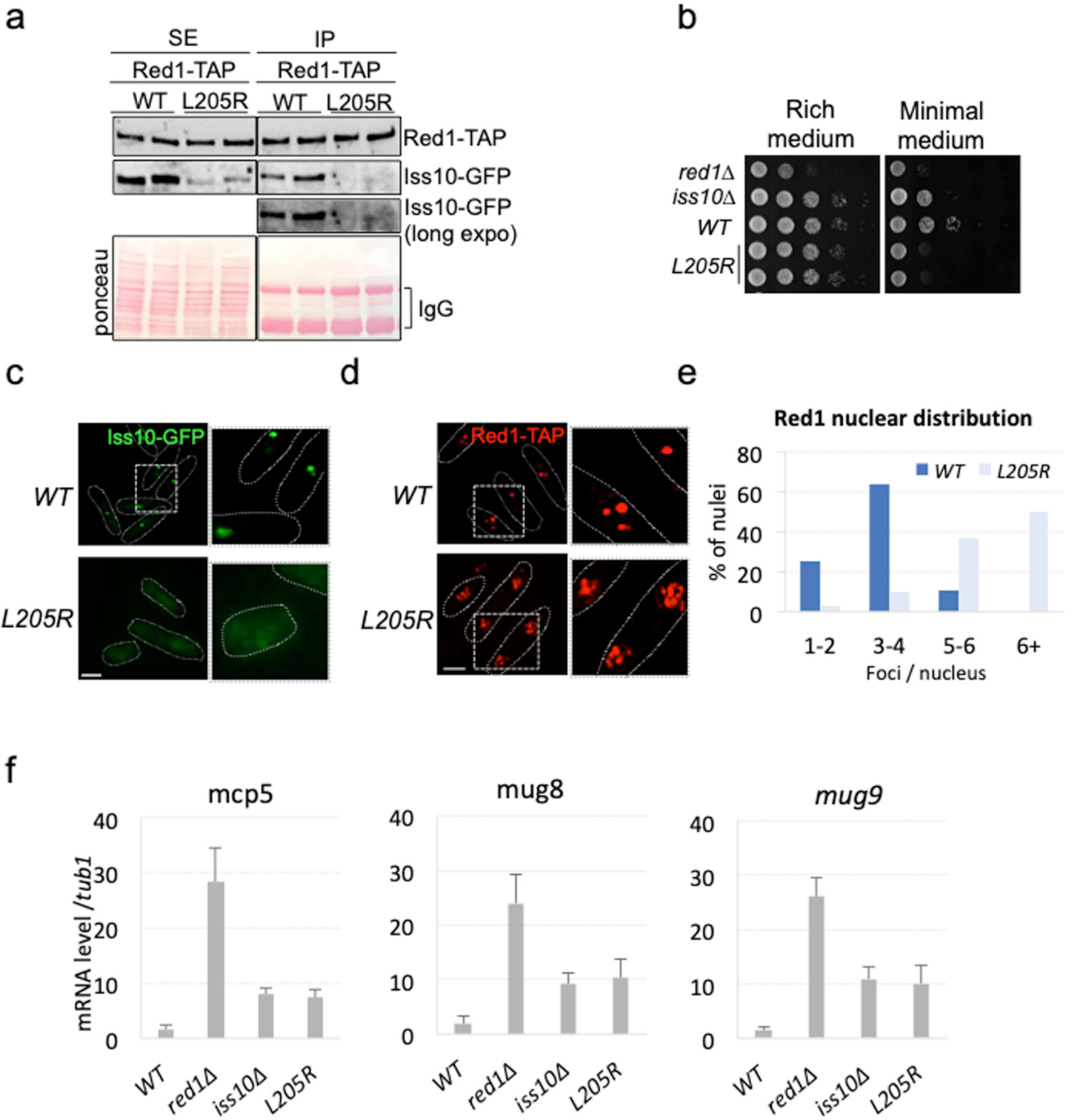
Red1^L205R^ mutation affects cell growth, degradation of DSR-containing mRNAs and clustering of Red1 foci. **(a)** Co-immunoprecipitation experiments, analyzed by western blot, showing that Red1^L205R^ mutation compromises Red1-Iss10 interaction *in vivo*. Cell extracts were prepared from exponentially growing cells expressing Red1-TAP and Iss10-GFP in liquid minimal medium. **(b)** Plating assays monitoring the growth of of *red1-L205R* compared to *WT, Iss10Δ* and *red1Δ* cells. Cells were grown on solid rich or minimal media for 5 days at 25°C. **(c)** Images of live fluorescence microscopy showing the localization Iss10-GFP in *WT* and *red1-L205R* cells. Scale bar = 5µm. **(d)** Images of immuno-fluorescence microscopy showing Red1-TAP localization in *WT* and *red1-L205R* cells. Scale bar = 5µm. **(e)** Graph showing the number of Red1 foci per nucleus in *WT* and *red1-L205R* cells. A minimum of 100 nuclei from two different biological isolates were counted. **(f)** Quantitative RT-PCR experiments showing the level of *mcp5, mug8* and *mug9,* three meiotic and DSR-containing mRNAs normalized to *tub1* mRNA in *WT, red1Δ, iss10Δ* and *red1-L205R* cells. Total RNAs were purified from cells growing exponentially in liquid minimal medium.

### Red1 interacts with Ars2 via its conserved N-terminal motif

To define the Red1 region interacting with Ars2, we again used Y2H assays. Analysis of the three consecutive Red1 fragments revealed that Ars2 specifically interacts with the Red1 N-terminus (1-140) (Supplementary Fig. 4a). These results were confirmed *in vitro* by size exclusion chromatography with purified proteins (Supplementary Fig. 4 b-e). Indeed, His-MBP-fusions of full-length Ars2 and Red1^1-140^ co-eluted in a single peak, distinct from the elution peaks of individual proteins (Supplementary Fig. 4b,c). To further define the minimal region of Red1 involved in the interaction with Ars2, several truncations of Red1^1-140^ were assayed, which eventually revealed that Red1^20-40^ is sufficient for Ars2 binding (Supplementary Fig. 4b-e). Interestingly, this Red1 fragment contains a short sequence motif “EDGEI”, which is significantly more conserved among several *Schizosaccharomyces* species than the rest of Red1^1-140^ (Supplementary Fig. 4f). To further characterize the interaction between Red1 and Ars2, we used isothermal titration calorimetry (ITC). Both Red1^1-54^ and Red1^20-40^ (with and without MBP tag) bound Ars2 with an equivalent dissociation constant (K_d_) of about 5 µM (Fig. 4a, Supplementary Fig. 5a,b), further supporting the hypothesis that Red1 region spanning residues 20-40 is a major determinant of the Ars2 binding.

**Figure 4.**
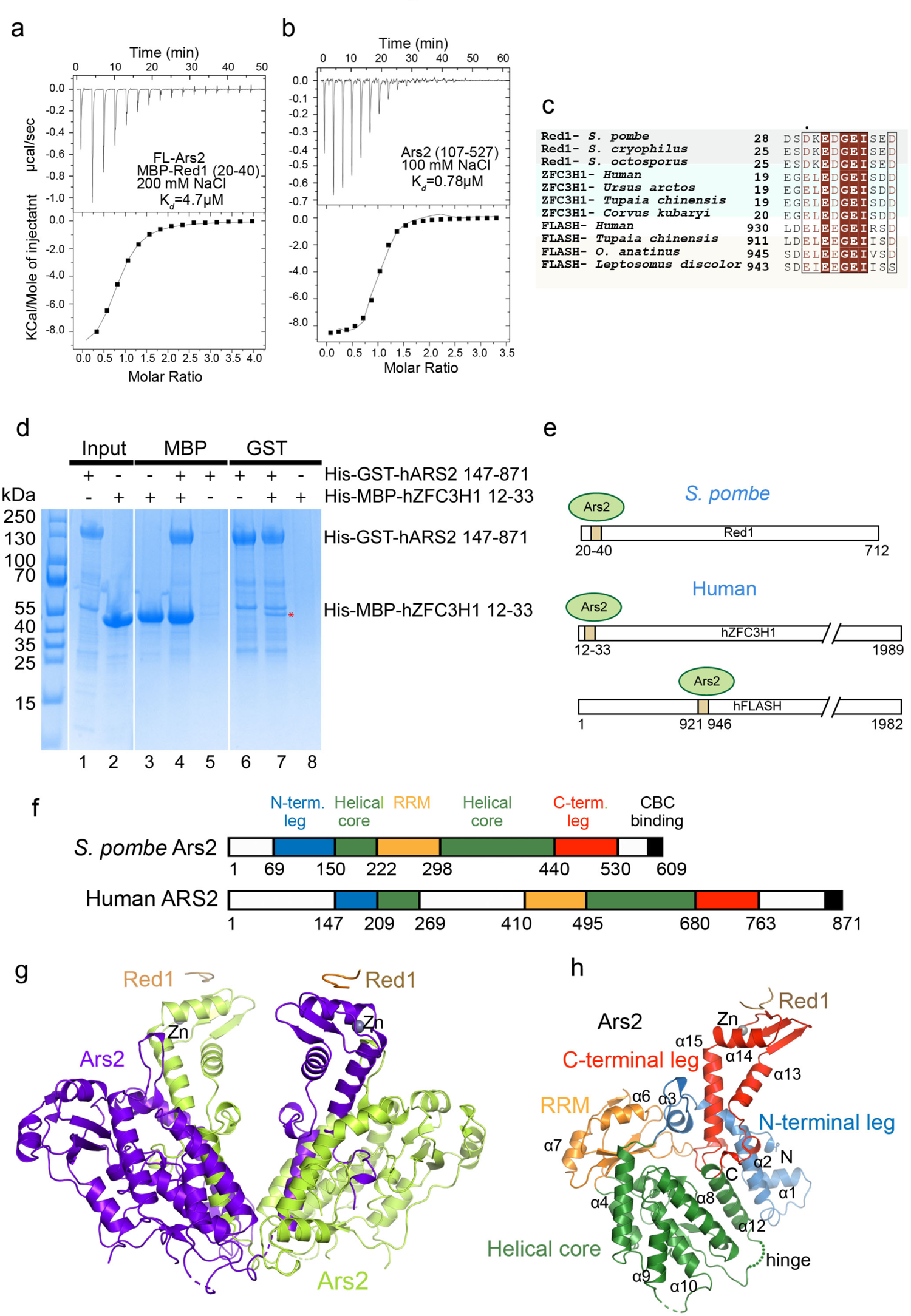
Red1 interacts directly with Ars2 via its EDGEI motif. **(a)** ITC measurement of the interaction affinity between full-length (FL) Ars2 and His-MBP-Red1^20-40^ in presence of 200mM NaCl. **(b)** ITC measurement of the interaction affinity between Ars2^107-527^ and His-MBP-Red1^20-40^ in presence of 100mM NaCl. **(c)** Sequence alignment of the Red1, hZFC3H1 and hFLASH proteins shown to interact with Ars2. Identical residues are in brown boxes. **(d)** MBP- and GST-tag pull-down experiments with His-GST-tagged human ARS2^147-871^ and His-MBP-tagged ZFC3H1^12-33^ showing, respectively, a mutual interaction (lanes 4 and 7). The proteins were first purified using Ni^2+^ resin (lanes 1,2) and then co-purified on either Amylose or GST resins. **(e)** A schematic diagram highlighting the conservation of the Ars2 binding motifs between *S. pombe* and human. **(f)** Schematic representation of the domain structure of *S. pombe* and human Ars2, as defined in this study and in Schulze *et al* ^32^. The CBC binding site was characterized in^39^. **(g)** Ribbon representation of the overall structure of the Ars2^68-183,206-531^ dimer. The C-terminal region (420-531) corresponding to the C-terminal leg is swapped between the protomers. The interacting peptide of Red1 is shown in brown. **(h)** Crystal structure of the Ars2 monomer with C-terminal leg not swapped in complex with Red1. The four defined domains are highlighted in different colours. Red1 peptide is in brown. The hinge region, allowing for the mobility of the C-terminal domain is shown. Annotated secondary-structure elements correspond to those shown in the alignment of Supplementary Fig. 6.

*S. pombe* Ars2 consists of 609 residues and in analogy to hARS2 is predicted to contain intrinsically disordered N- and C-termini and a central structured region. Based on deletion mutagenesis, limited proteolysis and sequence comparisons to hARS2, we generated the Ars2^107-527^ construct lacking both disordered termini, which turned to be more stable than the full-length protein and retained its interaction with Red1. The strength of the interaction depends on salt concentration, since when NaCl concentration was reduced from 200 to 100 mM, the K_d_ measured by ITC for Ars2^107-527^ and Red1^20-40^ increased from 5 to 0.8 µM (Fig. 4b). Together, these results demonstrate that the direct interaction between Red1 and Ars2 is mediated by a conserved EDGEI motif-containing segment spanning residues 20-40 of Red1 and the central structured core of Ars2.

### Conservation of the Ars2-Red1 binding interface in human

The ZFC3H1 subunit of the PAXT complex has been proposed to be the human counterpart of Red1^7^. ZFC3H1 (1989 residues) is much larger than Red1 and the two proteins differ significantly in sequence. However, there is a clear sequence match between the ZFC3H1 N-terminus and the Ars2 binding region of Red1 (Fig. 4c). This suggests that, analogous to the interaction of Ars2 with the MTREC complex via Red1, human ZFC3H1 might mediate the interaction of ARS2 with the PAXT complex. To confirm this hypothesis, we used pull-down experiments with GST-tagged human ARS2^147-871^ and MBP-tagged ZFC3H1^12-33^. We identified a direct interaction between ARS2 and ZFC3H1 mediated by the conserved ZFC3H1^12-33^ region (Fig. 4d, lanes 4 and 7). Importantly, this region contains the same EDGEI motif (Fig. 4c). Furthermore, it has previously been reported that human ARS2 interacts via a similar motif with FLASH, a large protein involved in histone mRNA biogenesis^32,35^. FLASH^931-943^ binds ARS2^147-871^ with a K_d_ of 0.5-5 µM depending on salt concentration^32^. The ED/EGEI motif mediated-interactions with ARS2 within the MTREC/PAXT complexes are thus conserved between human and *S. pombe*. Moreover, in human, at least two proteins (ZFC3H1 and FLASH) use this motif for direct binding to ARS2, most likely in a mutually exclusive way (Fig. 4e).

### Ars2-Red1 complex crystal structure determination

To gain molecular insights into the Ars2-Red1 interaction, we set out to determine the atomic structure of the complex by X-ray crystallography. First, we obtained crystals of the complex between Ars2^107-527^ and Red1^20-40^, that however only diffracted to 5 Å. Inspired by the recently available Alphafold model of *S. pombe* Ars2 (AF-O94326)^36^, we then extended both the N- and C-terminal limits of the Ars2^107-527^ construct and removed one predicted mobile loop (residues 184-205). This new construct was produced by co-expression of its two fragments - Ars2^68-183^ and Ars2^206-531^. Two separate elution peaks of Ars2 were obtained by size exclusion chromatography (Supplementary Fig 5c,d) and MALLS analysis identified them as a homodimer and a monomer, respectively (Supplementary Fig. 5e,f). Only the minor dimeric version of Ars2 in complex with Red1^20-40^ yielded single crystals, which diffracted to a 2.8 Å resolution. The structure was determined by molecular replacement using the Alphafold Ars2 model (AF-O94326) and refined to an *R*_free_ of 30.2% and an *R*_work_ of 24.5% (Supplementary Table 2).

The *S. pombe* Ars2 structure consists of four defined domains, including a helical core with an inserted RNA recognition motif (RRM) and is flanked by two extended, mostly helical structures. These extensions are referred to as the N- and C-terminal leg, in similarity to its human counterpart ^32^ (Fig. 4f-h). Ars2^68-183,206-531^ crystallised as a domain-swapped dimer, having the C-terminal leg segment exchanged between the two protomers (Fig. 4g). The hinge region, allowing the rotation of the C-terminal leg, corresponds to residues 420-421 (Supplementary Fig. 7a). The domain swap occurred likely already during the overexpression in bacteria as it could not be reproduced even with high concentrations (up to 40 mg/ml) of the monomeric Ars2. Since it is not known, whether such an Ars2 dimer has a physiological significance, in the following text, the monomeric Ars2 structure, without the domain swap, will be considered.

### The Ars2-Red1 interface

Red1 interacts with the Zn-finger domain of the Ars2 C-terminal leg, particularly with the surface formed by α14, β11 and β12 (Fig. 5a). In our Ars2-Red1 complex structure, electron density is visible for six out of the twenty Red1 peptide residues (Supplementary Fig. 7b). The peptide adopts a U-shape structure bent around the central Gly34 and establishes hydrophobic and charged contacts with Ars2, burying 300 Å^2^ of its surface. Red1 Gly34 and Ile36 together with aliphatic portions of Asp33 and Glu35 side chains pack against the Ars2 hydrophobic surface generated by Phe485, Phe490, Leu484, Leu486 and Lys473 (Fig. 5b). In addition, the three negatively charged residues Glu32, Asp33 and Glu35 make several hydrogen bonds and salt bridge interactions with Ars2 Lys483, Lys493 and, His494 (Fig. 5c). The electrostatic nature of these interactions is consistent with the variation of the K_d_ with salt concentration (Fig, 4a,b). Finally, Red1 Ile36 and Gly34 form main chain interactions with Ars2 Leu484 and Leu486 (Fig. 5c).

**Figure 5.**
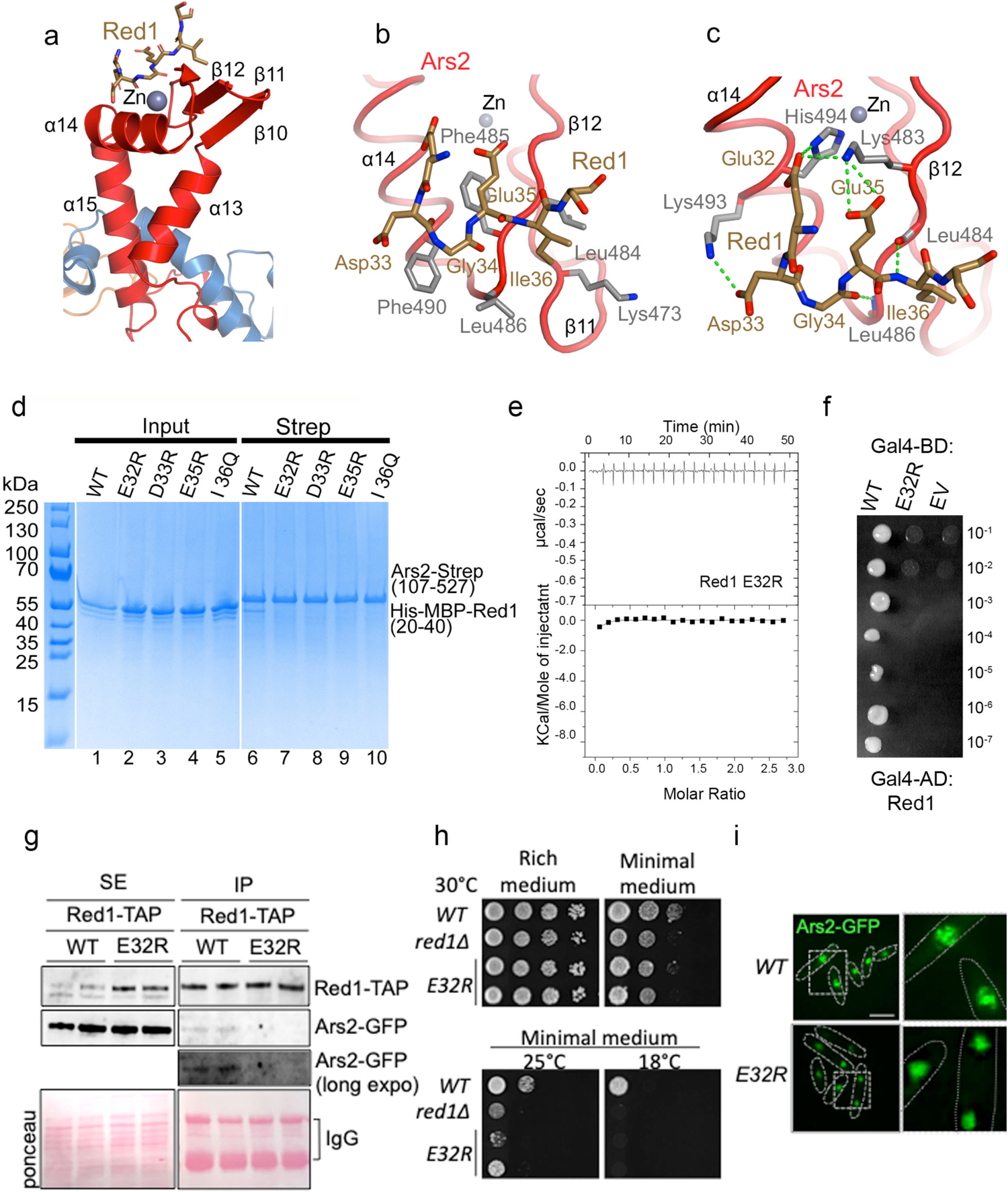
Red1-Ars2 interaction is necessary for proper cell proliferation and Ars2 nuclear localization into Red1 foci. **(a)** The Red1 peptide (in brown) interacts with the Zn-finger domain of the Ars2 C-terminal leg (in red). **(b)** Details of the interaction between the Red1 EDGEI motif and the Ars2 surface formed by α14, β11 and β12. Hydrophobic contacts between Red1 and the Ars2 residues are shown. **(c)** Residues of the Red1 EDGEI motif form several side and main chain hydrogen bonds and salt bridge interactions with the Ars2 C-terminus. **(d)** Strep-tag pull-down experiments with Strep-Ars2^107-527^ and His-MBP-Red1^20-40^ mutants indicated above the lanes. Ars2 and Red1 proteins were first purified using Ni^2+^ resin (lanes 1-5). Ars2 with either WT or mutant Red1 were then co-purified on Strep-tactin resin (lanes 6-10). All mutations abolished the interaction. **(e)** ITC measurement of the interaction affinity between Strep-Ars2^107-527^ and the His-MBP-Red1^20-40^ E32R mutant in presence of 100mM NaCl. **(f)** Essential role of Red1 Glu32 in the interaction with Ars2 as shown by Y2H. Growth was assayed on medium lacking histidine and with 30mM 3AT. **(g)** Co-immunoprecipitation experiments of Red1-TAP and Ars2-GFP in *WT* and *red1-E32R* cells, analysed by western blot. Soluble cell (SE) extracts were prepared from exponentially growing cells in minimal liquid medium. **(h)** Plating assays monitoring the growth of *WT, red1Δ and red1-E32R* cells on rich and minimal media at different temperatures. **(i)** Images of live cell fluorescence microscopy showing Ars2-GFP localization in *WT* and *red1-E32R* cells. Scale bar = 5µm.

In order to identify the key Ars2 and Red1 residues for stabilizing the interaction, we mutated Ars2 Lys483, which forms salt bridge interactions with Red1, as well as Phe490, which is central to the hydrophobic contacts between the two proteins (Fig. 5b,c). The integrity of the Ars2 mutants was verified by gel filtration, with an elution profile similar to the WT protein (data not shown). In Strep-tag pull down assays, both mutations (K483D, F490D) essentially disrupted the Red1 binding (Supplementary Fig. 7c, lanes 3,4). We also mutated key Red1 residues including Glu32, Asp33, Glu35 and Ile36 within the Red1 peptide (20-40). In pull-down assays with Strep-tagged Ars2^107-527^, mutations of all four residues abolished the binding (Fig. 5d, lane 7-10), confirming the importance of these residues for the interaction. In addition, the role of E32R was supported by ITC measurements, since no binding to Ars2^107-527^ was observed for the peptide containing the E32R mutation (Fig. 5e). Finally, we confirmed the essential role of E32R for the interaction with Ars2 in the context of full-length proteins using Y2H assays, which showed that Red1 E32R did not bind Ars2 (Fig. 5f). The structure-guided mutagenesis thus validated the key residues observed in the structure for the Red1-Ars2 interaction and identified the Red1 E32R mutation as a suitable candidate for functional analysis.

### *In vivo, red1-E32R* mutation affects Red1-Ars2 interaction, cell growth and Ars2 localization in nuclear foci

We then examined the function of Red1-Ars2 binding interface *in vivo.* We assessed the impact of *red1-E32R* point mutation on Red1-Ars2 interaction by co-immunoprecipitation experiments. Recombinant Red1-TAP, Red1-E32R-TAP and Ars2-GFP proteins were expressed instead of endogenous proteins and from respective endogenous promoters. Importantly, while Red1-TAP and Ars2-GFP interact, as expected, this interaction was lost in cells expressing Red1-E32R-TAP (Fig. 5g). Next, we conducted growth assays on different solid media and at different temperatures. The *red1Δ* cells experience a growth defect, which is exacerbated on minimal medium (Fig. 5h, Supplementary Fig. 3a,b) or at lower temperature (Supplementary Fig. 8a). A similar growth defect was visible for the *red1-E32R-TAP* mutant cells grown on minimal medium, suggesting that Red1-Ars2 interaction, depending on the growth conditions, can be important for the normal growth of the cell population. As MTREC degrades specific population of RNAs such as meiotic DSR-containing mRNAs and PROMPTs/CUTs ^7^, we asked whether the levels of these transcripts changed in *red1-E32R* mutant cells. However, no significant changes were observed for several meiotic mRNAs and PROMPTs/CUTs MTREC targets (Supplementary Fig. 8b,c). We also assessed the relevance of Red1-Ars2 interaction for their respective localization in nuclear foci. In cells expressing the Red1-E32R-TAP mutant protein, Ars2-GFP signal appeared to be more diffused and less concentrated in nuclear dots (Fig. 5i), while the localization of Red1-TAP in nuclear foci did not significantly change (Supplementary Fig. 8d,e). Thus, *in vivo*, Red1-Ars2 binding interface is critical for Red1-Ars2 interaction, for optimal cell growth and localization of Ars2 into nuclear foci where Red1 and other subunits of MTREC localize.

### Comparison of *S. pombe* and human Ars2

Compared to human ARS2, the mutual orientations of the domains in *S. pombe* Ars2 are considerably different. When the two proteins are aligned via their helical domains, large differences in positions of the N- and C-terminal legs and to a lesser extend of the RRM domain are apparent (Fig. 6a,b). Similar differences are also observed when compared to the *A. thaliana* SERRATE structure. The folds of individual domains, however, are rather well conserved between *S. pombe* and human (Supplementary Fig. 9a-d). Thanks to extending the N- and C-terminal limits of the construct used, the *S. pombe* Ars2 structure shows a more elaborate N-terminal leg, where the extreme N terminus (residues 69-80) packs against helices α1 and α2 on one side and interacts with C-terminal residues 527-530 on the other one (Supplementary Fig. 9e). The remaining part of the N-terminal leg resembles the one of human ARS2 (Supplementary Fig. 9a). The helical core Ars2 has a similar topology to its human counterpart (Supplementary Fig. 9b) and has the RRM domain inserted after helix α5. While the RRM domain β-sheet surfaces frequently carry exposed aromatic residues in the two RNP1 and RNP2 motifs that are involved in binding of ssRNA sequences^37^, the Ars2 RRM possesses only one Trp267 located on β3 (RNP1) (Supplementary Fig. 9c). In addition, the β-sheet and Trp267 are packed against α11 of the helical core, making it unavailable for interaction with RNA. The C-terminal leg can also be well superimposed onto the human C-terminus (Supplementary Fig. 9d). However, an important difference is the presence of a Zn atom coordinated by Cys476, Cys481, His455 and His494 in *S. pombe*, which is lacking in human ARS2, due to replacement of Cys481 by a serine (Fig. 6c). The Zn co-ordination is preserved in *A. thaliana* SERRATE^38^. The mutual orientation of Ars2 domains is unlikely to be affected by the C-terminal leg swap, as observed in our low resolution structure of Ars2^107-527^, where no domain swap occurs, while the Zn-finger region and the N-terminal helix α2 is partially disordered (Supplementary Fig. 9f).

**Figure 6.**
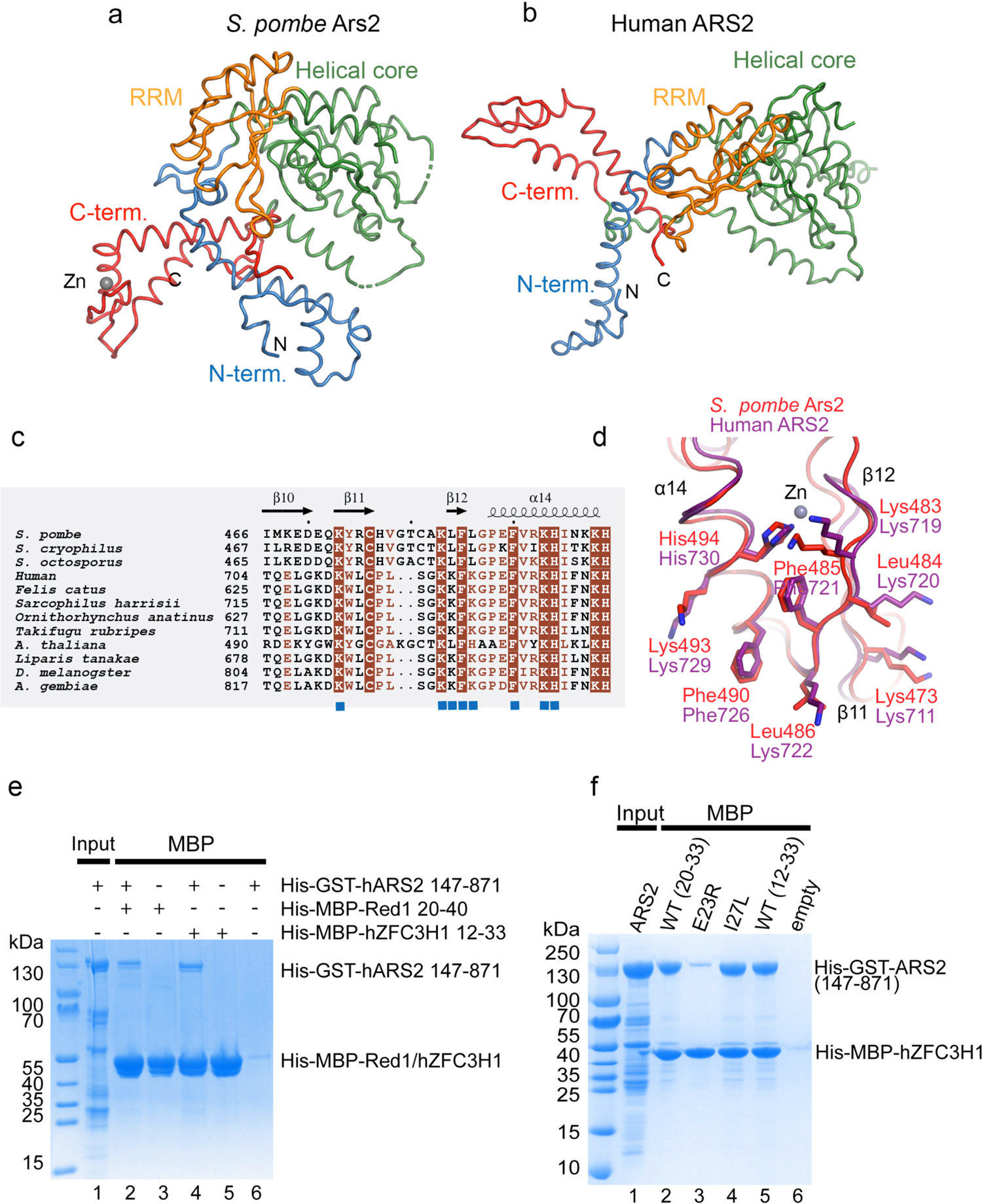
The Red1-binding surface in conserved in human ARS2. **(a)** Comparison of the *S. pombe* and **(b)** human Ars2 structures. The proteins were superimposed using the helical core domain (in green) with r.ms.d.−2.8 Å for 153 Cα atoms. **(c)** Sequence alignment of Ars2 proteins. Only the Zn-finger domain of the C-terminal leg is shown. Identical residues are in brown boxes. Blue squares indicate residues involved in the interaction with Red1. **(d)** The Red1 binding surface of *S. pombe* Ars2 is conserved in human ARS2. The secondary structure element labeling corresponds to *S. pombe* Ars2. **(e)** MBP-pull down experiment, showing that Human His-GST-ARS2^147-871^ can interact with both His-MBP-ZFC3H1^12-33^ and His-MBP-Red1^20-40^. **(f)** MBP-pull down experiment with human His-GST-ARS2 and His-MBP-ZFC3H1. ZFC3H1^20-33^ is sufficient for ARS2^147-871^ binding (lane 2). E23R mutation abolished the interaction (lane 3) while I27L retains a WT level of binding (lane 4).

### The Red1-Ars2 interface is conserved in human

The Ars2 residues interacting with Red1 are very well conserved across species, underlining the importance of this interaction (Fig. 6c). Analysis of the human and *A. thaliana* ARS2/SERRATE structures and the Alphafold model of *D. melanogaster* ARS2^36^, revealed that the Ars2 surface interacting with its ED/EGEI motif-containing partners is very well preserved (Fig. 6d, Supplementary Fig. 10a). With the exception of Leu484 and Leu486, the interacting residues remain identical (Fig. 6c,d). While in *A. thaliana* Leu486 is substituted by a histidine that might impact on the binding affinity, Leu484 and Leu486 are replaced by a lysine in some species that likely retains hydrophobic contacts with Ile33 (Fig. 6c, Supplementary Fig. 10a). To confirm the conservation of the Ars2 surface, we used MBP-pull down assays, where the GST-tagged human ARS2 (residues 147-871) bound equally well to MBP-Red1 or MBP-ZFC3H1 “EDGEI” motif-containing peptides (Fig. 6e). Accordingly, in the human system, a K719A, K722A and K734A triple mutant located at the FLASH/ZFC3H1 binding site (Supplementary Fig. 10a) abolished the hARS2-FLASH interaction^32^. To further characterise the ARS2-ZFC3H1 interaction in the human system, we generated His-MBP fusion of ZFC3H1^12–33^ containing the E23R mutation analogous to the above-described E32R mutation in Red1 (Fig. 5d-f). In an MBP-pull down assay, mutated ZFC3H1 did not interact with ARS2 any longer (Fig. 6f, lane 3). In addition to ZFC3H1, we identified short conserved sequences reminiscent to the ED/EGEI motif in other human proteins such as NCBP3 (41-EVEEGEL-47, 214-EAEEGEV-220). Since the putative NCBP3 motifs contain leucine or valine instead of Ile27 in ZFC3H1 (Ile36 in *S. pombe,* Fig. 5c), we wanted to verify whether the isoleucine residue is strictly required in the motif to be able to bind ARS2. I27L mutation however did not perturb the binding between ZFC3H1^12–33^ and ARS2 (Fig. 6f, lane 4). We could also show that a short peptide containing only 13 residues (ZFC3H1^20–33^) is sufficient for the interaction with ARS2 (Fig. 6f, lane 2). Together, these results show that the *S. pombe* Ars2 surface interacting with Red1 is well conserved among species and is used in human ARS2 to bind the PAXT complex subunit hZFC3H1 as well as FLASH.

### Dimerization of Red1 C-terminus

To shed light onto the observed Red1 propensity to self-associate, we assessed which part of Red1 is required for its eventual oligomerization. In Y2H assays, the C-terminal Red1 construct (residues 390-712) interacted strongly with full-length Red1 (Fig. 7a). In addition, the C-terminal region was sufficient for the interaction when tested against the three Red1 fragments (Fig. 7a). We used Alphafold^36^ to model the dimeric structure of the Red1 C-terminus. While most of the structure is predicted with low confidence, Alphafold predicts an antiparallel dimeric coiled-coil corresponding to residues 470-524 with very high per-residue confidence score (pLDDT) and the predicted aligned error plot indicates high confidence of the mutual positions of the two protomers (Fig 7b, Supplementary Fig. 11a). Equivalent predictions were obtained for *S. octosporus* and *S. cryophilus* Red1 (Supplementary Fig. 11a). The predicted structure revealed six complete heptad repeats engaged in classical coiled-coil packing (Fig. 7b-d). To test this prediction, a His-MBP-Red1^452-524^ fusion protein was analysed by size exclusion chromatography, where two elution peaks were observed (Supplementary Fig 12a,b). MALLS revealed that the two peaks correspond to dimeric and monomeric Red1^452-524^, respectively (Fig. 7e, Supplementary Fig. 12c). The dimeric Red1^452-524^ remained dimeric upon reinjection on the gel filtration column (Fig. 7e, Supplementary Fig. 12d). Importantly, an L481R, I485R double mutation, based on the structure model, is able to abolish the Red1 dimerization (Supplementary Fig. 12e,f). These results thus provide evidence for a coiled coil-mediated homo-dimerization of Red1. Given that Iss10, Ars2 and Mtl1 each interact with Red1 in a 1:1 or 2:2 stoichiometry, it is thus possible that these subunits are present in two copies within the MTREC complex (Fig. 7f). Importantly, Alphafold also predicts, with high accuracy, coiled coil-based homodimers for the PAXT and NEXT complex scaffolding proteins hZFC3H1 and ZCCHC8, respectively (Supplementary Fig. 13). The dimeric architecture might thus be a conserved feature among all these RNA targeting complexes.

**Figure 7.**
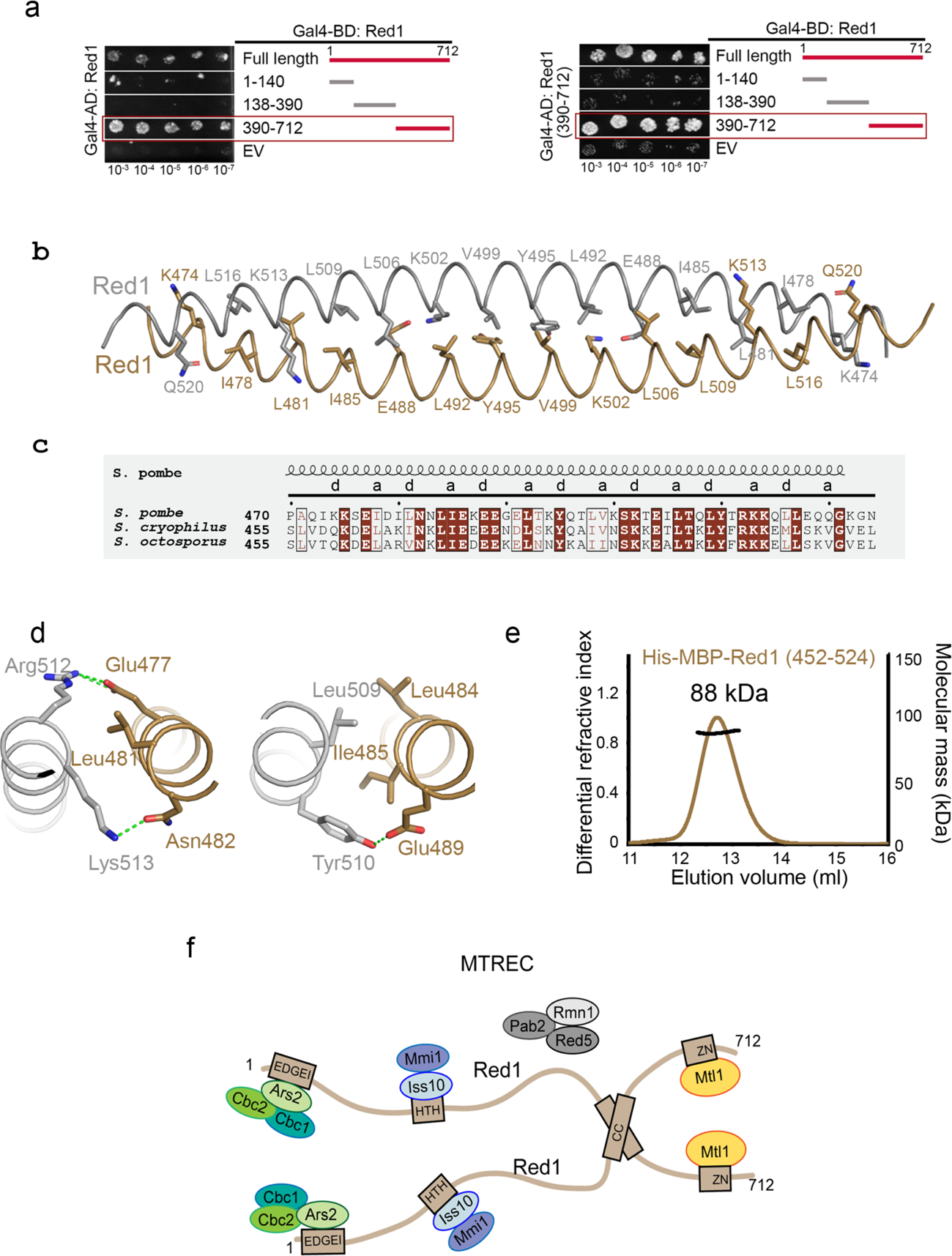
Red1 homo-dimerization. **(a)** Y2H mapping of the Red1 regions involved in its self-association. Growth was assayed on medium lacking histidine and with 30 and 50mM 3AT. EV-empty vector. FL-Red1 (upper panel) or Red1^390-712^ (lower panel) were used as a prey. **(b)** Ribbon representation of the Red1 dimeric coiled coil as modeled by Alphafold^36^. Residues in heptad positions indicated in **c** are shown as sticks. **(c)** Sequence alignment of Red1 proteins comparing different *Schizosaccharomyces* species. Only the sequence of the coiled-coil region is shown. Identical residues are in brown boxes, and conserved residues are highlighted in brown. Heptad positions are indicated. **(d)** Details of the Red1 dimeric coiled-coil interactions centred on Leu481 (right panel) and Ile485 (left panel). **(e)** Molecular mass determination of the His-MBP-Red^452-524^ by MALLS. The measured molecular mass of 88 kDa corresponds to a dimer. Calculated molecular mass of a monomer is 53 kDa. The sample was injected at 9 mg/ml. **(f)** Schematic model of the MTREC complex, summarising the structural and biochemical information obtained in this study and in^14^.

## Discussion

The *S. pombe* MTREC complex and its human counterpart PAXT target specific and aberrant nuclear transcripts for degradation by the nuclear RNA exosome^4, 5^. While the core RNA Mtl1/MTR4 helicase has been extensively analysed^15^, the exact roles of the remaining subunits remain unclear. In this study, we describe molecular details of the Red1 subunit interactions with Iss10 of the Iss10/Mmi1 sub-module and Ars2 of the cap-binding sub-module (Cbc1, Cb2, Ars2). In addition, we report a coiled-coil mediated homo-dimerization of Red1. Together with the recently reported crystal structure of the complex between Red1 and Mtl1^14^, these results provide molecular insights into the Red1 function as a scaffold of the MTREC complex, linking together individual sub-complexes. We also show that these MTREC features are conserved in the human analogue PAXT complex.

Our NMR structure of the minimal Red1-Iss10 interaction complex revealed that Red1 uses a HTH motif to bind the Iss10 N-terminal helices. We identified a Red1 L205R mutant that disrupts the Red1-Iss10 interaction and analysed its impact on MTREC functions in *S. pombe* cells. This mutation was sufficient to compromise the Red1-Iss10 binding *in vivo*, resulting in an Iss10 protein level decrease, indicating that the Red1 interaction is required for Iss10 stabilisation. Accordingly, the amount of Iss10 was previously shown to be drastically reduced in *red1Δ* cells^26^. Noticeably, *red1-L205R* cells show a growth defect similar to *red1Δ* cells on minimal medium and at lower temperatures that is stronger than in *iss10Δ* cells, but the reason for this is currently unclear.

Red1 is crucial for nuclear exosome foci formation^23^. Specifically, this function has been attributed to region 196–245^23^, which we now show forms a HTH domain required for Iss10 binding. Proper Red1 and exosome nuclear localization is also dependent on Iss10^6, 23^. Accordingly, we show that the *red1-L205R* mutation affects both Iss10 and Red1 localization. The interaction of Iss10 with Red1 HTH domain is required for Iss10 recruitment to Red1 nuclear foci, may promote the clustering of Red1 nuclear foci and enable recruitment of the nuclear exosome.

Expression of the Red1-L205R mutant instead of WT Red1 leads to accumulation of meiotic mRNAs that is comparable to *iss10Δ* mutation. This phenotype further raises when cells are grown in minimal medium, however, still remains lower than in *red1Δ* cells. The Iss10-Red1 interaction is thus dispensable for MTREC-based degradation of meiotic transcripts. Our Y2H assays also show that Iss10 interacts with Mmi1, known to directly bind to meiotic transcripts^21^, indicating that Iss10 links Red1 to Mmi1 and hence to target transcripts. The same Y2H interaction was recently reported in^14^. In *iss10Δ* cells, the interaction between Red1 and Mmi1 is severely compromised but not completely lost ^7, 26^. It is thus possible that while Iss10-mediated interaction of Mmi1 with Red1 is important for MTREC activity, particularly in sub-optimal conditions, Mmi1 might also be recruited into MTREC by alternative interactions as proposed in^14^. The fact that the impact of the Iss10-Red1 interaction varies according to the environmental growth conditions might be linked to the Iss10 regulation by TORC1^27^. Further analysis of the Mmi1 interaction with MTREC and Iss10 regulation by TORC1 will help explain the mechanism of MTREC-mediated degradation of meiotic transcripts in *S. pombe*.

We also report the crystal structure of the complex between *S. pombe* Ars2 and a Red1 N-terminal peptide. The structure of Ars2 reveals four structured domains that resemble their counterparts in hARS2 but their mutual orientations are very different, highlighting considerable plasticity of the Ars2 architecture in different species. While ssRNA binding has been reported for the hARS2 RRM domain^32^, the *S. pombe* Ars2 RRM is tightly packed against the helical core and its putative RNA binding residues are buried in the interface. hARS2 is involved in numerous interactions that are crucial for determination of the target transcript fate^30^. Only the interaction of the flexible, extreme ARS2 C-terminus with CBC has so far been characterised in detail^39^. In the first structure of the Ars2 core with a partner protein, we show that the Zn-finger domain of the Ars2 C-terminal leg interacts with the Red1 conserved EDGEI motif. We generated structure-based mutations in the Zn-finger domain and Red1 peptide that abolish the binding *in vitro* and showed that Red1 E32R mutation is sufficient to disrupt binding with Ars2 *in vivo*. The *Red1-E32R-TAP* mutant cells experience growth defect resembling the *red1Δ* cells, suggesting that Red1-Ars2 interaction is important for an optimal cellular growth. In addition, in mutant cells, Ars2 appears to be more diffused and less concentrated in nuclear dots. On the other hand, we did not observe any significant changes in the levels of several meiotic mRNAs and PROMPTs/CUTs known to be targets of MTREC, suggesting the interaction is not critical for degradation of these targets by MTREC.

We show that the Red1 EDGEI motif is conserved in human hZFC3H1, a subunit of the PAXT complex, as is the C-terminal leg interacting surface in human ARS2. Correspondingly, we show that hARS2 and hZFC3H1 interact and the E23R mutation in hZFC3H1 (equivalent to E32R in Red1) is sufficient to disrupt the complex. This is consistent with the demonstration that the hARS2 construct corresponding to the C-terminal leg (628-876) is a major determinant in triggering RNA decay of endogenous PROMPTs transcripts and in tethering assays^40^. Indeed, in tethering assays, the ARS2 activity was reduced upon siRNA silencing of hZFC3H1, but not of ZC3H18 nor ZCCHC8^40^. Moreover, either a K719A, K722A, K734A triple mutation, or the single K719A mutation, in the C-terminal leg of ARS2 inhibited its RNA decay triggering activity^40^. Here, we show that residues Lys719 and Lys722 are directly involved in recognizing the Red1/ZFC3H1 EDGEI motif. It seems highly plausible that this activity of human ARS2 thus depends on its EDGEI motif-mediated interaction with hZFC3H1. In agreement, the same ARS2 triple mutation prevented interaction of hARS2 with PAXT and NEXT components in cell-based assays^32^.

Human ARS2 has previously been shown to interact via its C-terminal leg with the EEGEI motif-containing FARB region of FLASH, a protein important for processing of replication dependent histone mRNAs, and this interaction was abolished by the K719A, K722A, K734A triple mutation^32^.

Our structure now provides an explanation for this interaction and implies a mutually exclusive binding of hARS2 to hZFC3H1/PAXT and FLASH. In addition to hZFC3H1 and FLASH, similar motifs are found in other important RNA regulators including NCBP3 (41-EVEEGEL-47, 214-EAEEGEV-220), ZC3H18 (189-EDDDGEI-195, 194-EIDDGEI-200) or SCAF1 (390-EIEEGEI-396).

Interestingly, NCBP3^1-182^ interacts with hARS2 in pull-down assays, which can be abrogated by the ARS2 C-terminal leg K719A, K722A, K734A triple mutation^32^. This suggests that NCBP3 is another RNA biogenesis factor engaging in a mutually exclusive interaction with the ARS2 C-terminal leg, although it remains to be shown whether or not the putative motifs mediate this. We speculate that the conserved binding surface on the ARS2 C-terminal leg accommodates multiple mutually exclusive interactions with competing partner proteins that exhibit the ARS2 binding motif ‘EE/DGEI/L’, and that this interaction is important for fate determination of targeted RNAs.

Finally, we also demonstrate the ability of Red1 to dimerize. Given that Iss10, Ars2 and Mtl1 can each bind to a single Red1, it is possible that the MTREC complex might exert its function as a dimeric assembly, where at least these subunits are present in pairs. This would be consistent with the reported dimerization of Mmi1^41^, which is a direct binder of meiotic transcripts in *S. pombe*^20–22, 42^.

Mmi1 is dimerized via its interaction with its dimeric partner Erh1 and this is important for meiotic transcript degradation by MTREC^41^. Importantly, Alphafold modeling predicts with high confidence dimeric coiled-coils also for hZFC3H1 of the PAXT and ZCCHC8 of the NEXT complex. Dimeric architectures might thus be a conserved feature of these RNA targeting complexes.

In conclusion, our structural and *in-vivo* analyses of the MTREC/PAXT scaffolding proteins provide important insights towards elucidation of the molecular architecture and function of these essential RNA regulators.

## Methods

### Yeast two-hybrid (Y2H) Assay

Genes coding for MTREC subunits and their variants were cloned into pDONR221 vector using BP clonase recombination (Life Technologies). The inserts were then introduced into Y2H destination vectors (pDEST32 for bait and pDEST22 for prey) using Gateway cloning system (ProQuest-Life Technologies). Bait and prey plasmids were co-transformed into MaV203 yeast strain and plated on Nitrogen-base (NB) agar medium containing 2% glucose, Histidine, Adenine and Uracil and incubated for 48h at 30°C. Co-transformed colonies were replica-plated on the same medium and incubated for two more days at 30°C. Co-transformants were then assayed for *HIS3* reporter gene activation, by plating them on NB agar medium (2% glucose, Ade, Ura) supplemented with different concentrations (5-50mM) of 3-aminotriazole (3AT; Sigma-Aldrich) histidine inhibitor. Plates were incubated at 30°C and growth was monitored. Control plasmids from ProQuest™ two-hybrid system (Life Technologies). Co-transformation with empty prey vector was used as negative control. In the initial screen, Krev/RalGDS interaction was used as a positive control and RalGDS prey vector was also used as a mock control.

### Pull-down Assay

All variants of Red1 were cloned as His-MBP fusions into the pETM41 vector. Iss10 constructs were cloned as N-terminal Strep-tag fusions into pACYCDuet (Novagen). Combinations of Red1 and Iss10 constructs were co-expressed in *E. coli* BL21Star (DE3, Invitrogen). Following cell disruption, the Red1/Iss10 proteins-containing supernatants were loaded onto a Strep-Tactin XT resin (IBA) that was then extensively washed with a buffer containing 20mM Tris pH 8, 100mM NaCl and 5mM β-mercaptoethanol. Bound proteins were eluted by addition of 50mM of D-Biotin, and analyzed on 15% SDS-PAGE. For analysis of the impact of Red1 mutants on the interaction with Iss10, half of each culture was used for Strep-tag pull-down assays, as described above, and the other half for MBP pull-down assays. Supernatants were applied on Amylose resin (NEB) that was extensively washed and the bound proteins were eluted with addition on 10mM maltose.

Ars2-Red1 complexes were analysed in Strep-tag pull-down assays. Ars2^107–527^ and its variants were cloned to contain an N-terminal His-MBP- and C-terminal Strep-tag into pETM41. His-MBP fusions of Red1 proteins and Ars2 variants were expressed individually in *E. coli* BL21Star (DE3) and purified using a Ni^2+^ chelating Sepharose (GE Healthcare). Strep-Ars2 proteins were applied on Strep-Tactin XT columns and washed extensively. Equal amounts of Ni-purified His-MBP Red1 proteins were added. Columns were extensively washed with a buffer containing 20mM Tris pH 8 and 100mM NaCl and bound proteins were eluted with addition of 50mM of D-Biotin and analyzed by 12-15% SDS-PAGE.

For analysis of the interaction between human ARS2 and ZFC3H1, hARS2^147-871^ was cloned into pETM30 as a His-GST fusion and variants of ZFC3H1 as His-MBP fusions into pETM41. The proteins were expressed individually in *E. coli* BL21Star (DE3) and purified on a Ni^2+^ chelating Sepharose (GE Healthcare). His-GST-ARS2 was applied onto a GST resin (GE Healthcare) and His-MBP-ZFC3H1 onto an Amylose resin (NEB). After extensive washing, His-GST-ARS2 was added onto the ZFC3H1-bound MBP column while His-MBP-ZFC3H1 was added onto the ARS2-bound GST resin. Columns were washed with a buffer containing 20mM Tris pH 8 and 100mM NaCl, the proteins were eluted by addition of 10mM reduced glutathione or 10mM maltose, respectively, and analysed on 15% SDS-PAGE.

### Isothermal Titration Calorimetry (ITC)

ITC experiments were performed at 25°C using an ITC200 microcalorimeter (MicroCal). Experiments included one 0.5µl injection and 15-20 injections of 1.5-2µL of 0.4-0.8 mM His-MBP-Red1 (1-53, 20-40 or 20-40 E32R) or the Red1^20-40^ peptide into the sample cell that contained 30-40 µM Ars2 (FL, 58-609 or 107-527) in 20 mM Tris pH 8.0 and 100-200 mM NaCl. The initial data point was deleted from the data sets. Binding isotherms were fitted with a one-site binding model by nonlinear regression using the Origin software, version 7.0 (MicroCal).

### Multi Angle Laser Light Scattering (MALLS)

Size exclusion chromatography (SEC)-Light scattering (LS) experiments were conducted at 4 °C on an HPLC chromatography system consisting of a degasser DGU-20AD, a LC-20AD pump, an autosampler SIL20-ACHT, a communication interface CBM-20A and a UV-Vis detector SPD-M20A (Schimadzu, Kyoto, Japan), a column oven XL-Therm (WynSep, Sainte Foy d’Aigrefeuille, France), a static light scattering miniDawn Treos, a dynamic light scattering DynaPro Nanostar and a refractive index Optilab rEX detectors (Wyatt, Santa-Barbara, USA). The analysis was carried out with the software ASTRA, v5.4.3.20 (Wyatt, Santa-Barbara, USA). Samples of 20 µL were injected at 0.5 mL min-1 on a Superdex 200 10/300 GL (GE Heathcare), equilibrated with 20mM Tris-HCl pH8, 100mM NaCl, 5mM β-mercaptoethanol. Bovine Serum Albumin, at 2 mg.mL-1, in PBS buffer was injected as a control.

### Protein Expression and Purification

His-MBP-Red1^192-236^ and Strep-Iss10^1-45^ were co-expressed in *E. coli* BL21Star cells. After affinity chromatography using the Ni^2+^ chelating resin (GE Healthcare) in a buffer containing 20 mM Tris pH 8, 200 mM NaCl and 5 mM beta-mercaptoethanol, the His-MBP-tag on Red1 was cleaved off by TEV protease. The complex was further purified on Strep-Tactin XT resin and Superdex 200 size exclusion chromatography.

Ars2^68-183^ was cloned into pProEXHTb (Invitrogen) as a His-tag fusion and Ars2^206-531^ into pRSFduet vector (Novagen) as a Strep-tag fusion. The two fragments were co-expressed in *E. coli* BL21Star (DE3, Invitrogen) and the complex purified by affinity chromatography using the chelating Ni^2+^ Sepharose (GE Healthcare). After His-tag cleavage with the TEV protease, the two fragments were co-purified on Strep-Tactin XT resin (IBA). The final purification step was a Superdex 200 size-exclusion chromatography using a buffer containing 20mM Tris pH 8, 100mM NaCl and 5mM β-mercaptoethanol.

### NMR Spectroscopy

NMR spectra were recorded at 298 K using a Bruker Neo spectrometer at 700 MHz or 800 MHz, equipped with a standard triple resonance gradient probe or cryoprobe, respectively. Bruker TopSpin versions 4.0 (Bruker BioSpin) was used to collect data. NMR data were processed with NMR Pipe/Draw^43^ and analysed with Sparky 3 (T.D. Goddard and D.G. Kneller, University of California).

### Chemical shift assignment

Backbone ^1^H^N^, ^1^H^α^, ^13^C^α^, ^13^C^β^, ^13^C’ and ^15^N^H^ chemical shifts for the minimal Iss10-Red1 heterodimer were assigned from a sample of 500 µM [^13^C,^15^N]Red1^192-236^-Iss10^1-45^ using 2D ^1^H,^15^N-HSQC, 3D HNCO, 3D HNCACO, 3D HNCA, 3D HNCACB, 3D CBCACONH, 3D HNHA and 3D HACACONH spectra. Using the same sample, aliphatic side chain protons were assigned based on 3D H(C)(CO)NH-TOCSY (40 ms mixing time), and 3D (H)C(CO)NH-TOCSY (40 ms mixing time) spectra. This was complemented by 2D ^1^H,^13^C-HSQC, 3D (H)CCH-TOCSY (16 ms mixing time), and 3D (H)CCH-TOCSY (16 ms mixing time) spectra collected on the sample after exchange to 99% (v/v) D_2_O. Assignment of sidechain asparagine δ2 amdies and glutamine ε2 amides used a 3D ^15^N-HSQC-NOESY (120 ms mixing time). Stereospecific ^1^H-^13^C assignment of leucine and valine methyl groups used a 2D constant-time ^13^C-HSQC spectrum with a 150 µM sample of 10% ^13^C-labelled Red1^192-236^-Iss10^1-45^ in 99 % D_2_O. Aromatic ^1^H chemical shifts were assigned using the same sample with 2D TOCSY (60 ms mixing time) and 2D DQF-COSY spectra.

### NMR structure calculation

Structure ensembles were calculated using Aria 2.3/CNS1.2^44,45^. The ^1^H distances were obtained using NOE crosspeaks from a sample of 500 µM [^13^C,^15^N]Red1^192-236^-Iss10^1-45^ using a 3D ^15^N-HSQC-NOESY (120 ms mixing time) and 3D ^13^C-HSQC-NOESY (120 ms mixing time). Distance restraints were also obtained from the 150 µM sample of Red1^192-236^-Iss10^1-45^ in 99 % D_2_O from a 2D ^1^H,^1^H-NOESY (120 ms mixing time) spectrum. Protein dihedral angles were obtained by using TALOS-N^46^ and SideR^47,48^. Hydrogen bond restraints (two per hydrogen bond) were introduced following an initial structural calculation and include only protected amides in the centre of helices. Starting at iteration four in the structure calculations, residual dipolar coupling (RDC) values were included by measuring interleaved spin state-selective TROSY experiments. The aligned sample contained 17 mg/mL Pf1 phage (Asla Biotech). RDC-based intervector projection angle restraints used D_a_ and R values of 8 and 0.33, respectively. Final ensembles were refined in explicit water and consisted of the 20 lowest energy structures from a total of 100 calculated models. Complete refinement statistics are presented in Supplementary Table 1.

### X-ray Structure Determination

Pure Ars2^68–183,206-531^ was concentrated to 8 mg ml^-1^ in a buffer containing 20mM Tris pH 8, 100mM NaCl, and 5mM β-mercaptoethanol using Amicon ULTRA concentrator (Millipore). The pure protein was supplemented with a three-fold molar excess of Red1 peptide (20-KNEEDESNDSDKEDGEISEDD-40) (PSL GMBH). The complex was crystallized using hanging drop vapor diffusion method at 5°C. The best diffracting grew within three days in a solution containing 200mM Ammonium Formate and 20% (w/v) PEG3350. For data collection at 100 K, crystals were snap-frozen in liquid nitrogen with solution containing mother liquor and 30% glycerol.

Crystals of the *S. pombe* Ars2^68-183,206-531^ - Red1^20-40^ complex belong to the space group *P*2_1_ with the unit cell dimensions *a* = 64.670, *b* = 128.281, *c* = 89.3 and β =108°. The asymmetric unit contains a 2:2 Ars2/Red1 heterotetramer with a swapped C-terminus and has a solvent content of 63%. A complete native dataset was collected to a 3.1 Å resolution, partially extending to 2.8 Å, on the ESRF beamline ID30A-3. The data were processed using autoPROC^49^. Phases were obtained by molecular replacement using PHASER^50^ with an adjusted Alphafold model of *S. pombe* Ars2 (AF-O94326) as a search model. The initial map was improved using the prime-and-switch density modification option of RESOLVE^51^. After manual model rebuilding with COOT^52^, the structure was refined using Refmac5^53^ with NCS restraints to a final *R*_free_ of 30.2% and an *R*_work_ of 24.5% (Table 1) with 99.8% residues in allowed (94% in favored) regions of the Ramachandran plot, as analyzed by MOLPROBITY^54^. A representative part of the 2*F*_o_ − *F*_c_ and omit *F*_o_ − *F*_c_ electron density omit maps calculated using the refined model with and without the Red1 peptide, respectively, are shown in Supplementary Figure 7.

### Data deposition

The structure ensemble of the Red1^192-236^-Iss10^1-45^ heterodimer has been deposited at the Protein Data Bank (http://www.ebi.ac.uk/pdbe/) with accession ID 7QUU. Corresponding chemical shift assignments have been deposited in the Biological Magnetic Resonance Data Bank (http://bmrb.wisc.edu/) under BMRB accession number 34702. The atomic coordinates and structure factors of the *S. pombe* Ars2-Red1 complex have been deposited under the PDB accession codes XXXX.

### Standard Yeast Genetics

Strains were constructed by standard genetic techniques (Moreno et al., 1991), cultured at 30°C and under agitation at 220 rpm, in YEA (Rich medium) or Edinburgh minimal medium with appropriate supplements (Minimal medium). Genotypes of strains used in this study are listed in Supplementary Table 3 Red1-TAP, Ars2-GFP and Iss10-GFP *S. pombe* cells were obtained using the PCR-based gene targeting method^55^ Positive transformants were selected by growth on YEA medium containing the appropriate antibiotic and confirmed by genomic PCR. Cells expressing point mutants of Red1 protein were generated in two steps from parental cells expressing *red1-TAP*. First, *ura4* gene was integrated into *red1* locus in order to generate *red1-Δ30-207.* Positive transformants were selected by growth on medium deprived of uracil and validated by genomic PCR. Then, *red1Δ 30-207::ura4+* cells were transformed with synthesized DNA fragments of *red1* (GeneArt) containing either the E32R or L205R mutations. Positive transformants were selected by growth on YEA-5FOA and validated by genomic PCR and DNA sequencing. Oligos used for the different strain generation are listed in Supplementary Table 4.

Cells co-expressing one of the different versions of Red1-TAP (WT, E32R or L205R) together with Ars2-GFP or Iss10-GFP were obtained by crossing cells with appropriate genotypes using random spore analysis and selection on the appropriate media. *S. pombe red1-TAP, iss10Δ* cells were obtained by crossing *iss10Δ* strain with *red1-TAP* strain and selected on the appropriate medium after random spore analysis.

### Cell Growth Spotting Assay

Spotting assays were done from 10^7^ cells of exponentially growing cultures (OD_600nm_ < 2) in minimal media. A 1/10th serial dilution of cells was dropped on rich or minimal media and grown at different temperatures (30, 25 and 18°C). Pictures of the plates were acquired (Syngene bioimaging camera) after 3 days for the plate grown at 30°C, 5 days for the plates grown at 25°C and 10 days of grown at 18°C.

### Live Cell Imaging

Red1-TAP (WT), Red1-TAP-E32R (E32R) and Red1-TAP-L205R (L205R) *S. pombe* cells co-expressing Ars2-GFP or Iss10-GFP were grown on liquid minimal medium. Live microscopy analysis was performed when cells reached an OD_600_ of 0.5-0.8. Images were acquired with a Zeiss Apotome microscope (Carl Zeiss MicroImaging) and using a 63X oil immersion objective with a numerical aperture of 1.4 (WD 190, DICIII and GFP filters). Raw images were analyzed using the AxioVision software (Carl Zeiss MicroImaging), and processed using ImageJ.

### Immunofluorescent Microscopy

The same strains used for the live cell imaging were used to assess the Red1-TAP localization by detecting its TAP tag using immunofluorescence experiments. Briefly, 20ml of *S. pombe* cell cultures in logarithmic phase (OD_600 =_ 0.5-0.8) and grown in minimal medium were fixed with 3.8% paraformaldehyde at room temperature for 30 min, then washed with 10ml PEM (100mM Pipes, 1mM EGTA, 1mM MgSO4 pH6.9). The cells were then transferred to 1.5ml Eppendorf tubes with 1ml of PEM and washed 2 more times. Cells were resuspended in 1ml PEMS (PEM, 1M D-Sorbitol) with 0.25mg Novozym (Sigma) and 0.25mg Zymolyase (Sigma) and incubated at RT (ideal @30°C) until > 90% cells are digested, then washed 3 times in 1ml PEMS and spin 30 seconds at 500g. Next, the cells were permeabilized 2 min with PEMT (PEM 1% Triton X-100), washed once with 1ml of PEM and then blocked with PEMBAL (PEM, 1% BSA, 100mM L-Lysine) for 30 min. Detection of Red1-TAP was obtained by incubating the cells for 2 hours at RT with the anti-TAP primary antibody (Thermo scientific #CAB1001) (1:100) in PEMBAL on a rotating wheel, the cells were washed 3 times 10 min with PEMBAL. The cells were then incubated over night at 4°C on rotating wheel (or 2 h at RT) with a secondary antibody coupled to a fluorescent dye (DyLight® 549 (VECTOR lab # DI 1549) (1:400) dilution in PEMBAL, cells are washed 3 times 10 with 1ml of PEMBAL then resuspend in 100µl PEMBAL and mounted on poly L-lysine coated coverslips. Fluorescence microscopy imaging were done with a Zeiss Apotome microscope (Carl Zeiss MicroImaging) and using a 63X oil immersion objective with a numerical aperture of 1.4 (WD 190, DICIII and GFP filters). Raw images were acquired using the AxioVision software (Carl Zeiss MicroImaging), and processed using ImageJ.

### Protein Co-immunoprecipitation

20ml of *S. pombe* cells were grown to mid-log phase in minimal medium at 30 °C, harvested, and flash-frozen in liquid nitrogen prior to protein extract preparations. Extracts prepared from cells expressing epitope tagged proteins under the control of native gene promoters. Red1-TAP WT or mutated versions (E32R, L205R) were used for immunoprecipitations to assess the interaction between Red1-TAP (WT or E32R point mutant) with Ars2-GFP, and between Red-TAP (WT and L205R point mutant) and Iss10-GFP. Total protein extracts were prepared from cells lysated mechanically using glass beads (2x 60sec) in IP-Lysis Buffer (50mM HEPES-NaOH pH=7.5, 5mM MgCl_2_, 20mM β-glycerophosphate, 10% glycerol, 1mM EDTA, 1mM EGTA, 50mM NaF, 0.1mM Na_3_Vo_4_, 0.2% NP-40, 150mM NaCl, 1µg LABP, 1mM Benzamidine, 1mM PMSF). Soluble extract cleared of cellular debris was incubated with IgG sephorose (GE Healthcare) for 2h at 4°C on a rotating wheel. Afterwards, beads were washed three times with the Lysis Buffer. Protein elution was performed using 2x Leammli buffer (50mM Tris-HCl, pH 6.8, 10% ß-mercaptoethanol, 1% SDS, 10% glycerol and 0.1% bromophenol blue) and by heating the samples 10 min at 70°C. The IP samples and 5% of the input were then separated by SDS-PAGE, transferred to nitrocellulose membranes and analyzed using appropriate primary (anti-TAP-Thermo scientific #CAB 1001, anti-GFP-Sigma # 11814460001) and secondary antibodies (HRP goat anti-Mousse-Dako #P0447, HRP goat anti-Rabbit-Dako #0448). Antibodies were diluted in TBS (0.1% Tween20-1% fat dry milk; 1:1000 for the primary and 1:10000 for the secondary), and membranes were washed in TBS-0.1% Tween20.

Enhanced chemiluminescence (ECL) detection was performed using reagents from Biorad (Clarity Western ECL kit) and revealed using ChemiDocMPSystem (BioRad) or the Fusion FX (Vilber) camera. Signals were quantified with the corresponding software, the imageLab software (Biorad) or the Fusion-Capt software (Vilber Lourmat).

### Quantitative RT-PCR

Total RNA was extracted from *S. pombe* cells using hot phenol-acid. Briefly, 25ml of logarithmic phase (OD_600 =_ 0.5-0.8) cells cultivated in minimal medium were collected by centrifugation 5 min at 1500g. Cells were washed with 1ml of PBS, then lysed by the addition 750 µl of TES (10mM Tris pH 7.5; 10mM EDTA pH 8; 0.5% SDS) and 750 µl cold acidic phenol-chloroform (Sigma P-1944), incubated at 65°C for 15 min, and vortexed 10 sec every 3 min during the heat incubation. Samples were then placed on ice for 1 min, vortexed 20 sec and centrifuged for 15 min at 12000g at 4°C. A total of 700 µl of the upper phase was recovered into a phase-lock (heavy) tubes (Eppendorf) and a second phenolic treatment was performed by the addition of 700 µl of acidic phenol-chloroform and mixing thoroughly by inverting the tubes. Samples were then centrifuged 5 min at 12000g at 4°C. The upper phase was transferred to new phase-lock tubes, and 700 µl of chloroform (Sigma) was added to the samples, mixed thoroughly again by inverting the tubes, and centrifuged 5 min at 12000g at 4°C. The aqueous phase (500 µl) was carefully transferred to a 2ml Eppendorf tubes containing 1.5ml of 100% cold EtOH (−20°C) and 50µl of 3 M NaAc pH5.2, the mix was vortexed and incubated 15 min at −80°C then centrifuged for 10 min at 12000g, 4°C. The pellet was washed in 75% (v/v) EtOH and centrifuged at 7500g for 5 min, 4°C. The pellet was then air-dried, dissolved in 100µl RNase-free water, incubated at 60°C for 10 min and stored at −20°C. RNA concentration and the A260/280 ration were determined using a NanoDrop ND-1000 UV spectrophotometer. RNA integrity was evaluated by electrophoresis on 1 % agarose gel using the GelGreen Nucleic Acid Stain (Biotium) dye following the manufacturer’s instructions.

Reverse transcription and PCR were performed on 2µg of purified RNA using the Transcriptor Reverse Transcriptase (Roche). RNA was first subjected to DNAse (Roche) treatment in the presence of 5X DNAse buffer (100mM Tris pH 8, 10mM MgCl_2_), 100mM DTT (Invitrogen) and RNase inhibitor (Thermo Scientific) for 20 min at 37°C. Next 0.7µl of 50mM EDTA were added to the RNA-DNAse mixture and incubated 10 min at 70°C. RNA was then hybridized with 50µM of random hexamer primer (Invitrogen) in the presence of 50mM MgCl_2_ for 10 min at 65°C. The cDNA synthesis was performed by adding 10mM dNTP (Thermo Scientific), RNase Inhibitor (Thermo Scientific), RT buffer (Roche) and Reverse Transcriptase (Roche) and subsequent incubation for 10 min at 25°C, 40 min at 55°C and 5 min at 85°C, according to the manufacturer guidelines.

qPCR was performed on a LightCycler480 machine (Roche). 4 µl of the diluted (1:4) cDNAs were used with 0.1µM of each primer (forward and reverse) and 2X MESA Blue qPCR mix (Eurogentec) in a 20µl final qPCR reaction. DNA amplification was done using the following program: 15 min incubation at 95°C, followed by 40 cycles of 95°C for 15 sec, 60°C for 15 sec and 70°C for 15 sec. Oligos used for the qRT-PCRs are listed in Supplementary Table 4.

## Acknowledgements

JK was funded by the CNRS ATIP-Avenir program. AV, SC and JK were funded by ANR MTREC (ANR-21-CE11-0021-01) and ANR RNAGermSilence (ANR-13-BSV2-0012) to AV. Ariadna B. Juarez-Martinez was supported by the Labex GRAL (Grenoble Alliance for Integrated Structural Cell Biology) (ANR-10-LABX-49-01) and the People Programme (Marie Curie Actions) of the European Union’s Seventh Framework Programme (FP7/2007-2013) under REA grant agreement PCOFUND-GA-2013-609102, through the PRESTIGE programme coordinated by Campus France. Financial support from the Centre Nationale de la Recherche Scientifique (IR-RMN-THC Fr3050) is gratefully acknowledged This work used the platforms of the Grenoble Instruct-ERIC center (ISBG; UAR 3518 CNRS-CEA-UGA-EMBL) within the Grenoble Partnership for Structural Biology (PSB), supported by FRISBI (ANR-10-INBS-0005-02) and GRAL, financed within the University Grenoble Alpes graduate school (Ecoles Universitaires de Recherche) CBH-EUR-GS (ANR-17-EURE-0003). We thank Caroline Mas for assistance with ITC, Luca Signor for mass spectrometry analysis, Aline Le Roy for assistance and access to the Protein Analysis On Line (PAOL) platform”, the staff of the ESRF-EMBL (European Synchrotron Radiation Facility-European Molecular Biology Laboratory) Joint Structural Biology Group, particularly Andrew McCarthy and Matthew Bowler, for access to and help with the ESRF beamlines as well as the IAB microscopy platform (MicroCell). We thank the EMBL high-throughput crystallization facility (HTX). We thank Axelle Grélard, Estelle Morvan, and the structural biology platform at the Institut Européen de Chimie et Biologie (UMS 3033) for access to NMR spectrometers, equipment, and technical assistance.We thank Rachel Fitzgerald for help with initial validation of Red1-Ars2 interaction and Wiebke Schulze for providing hARS2 expression plasmids.

