## Supplementary Figures and Tables for "Structural analysis of Red1 as a conserved scaffold of the RNA-targeting MTREC/PAXT complex"

Contains 13 Supplementary Figures, 4 Supplementary Tables and Supplementary references.

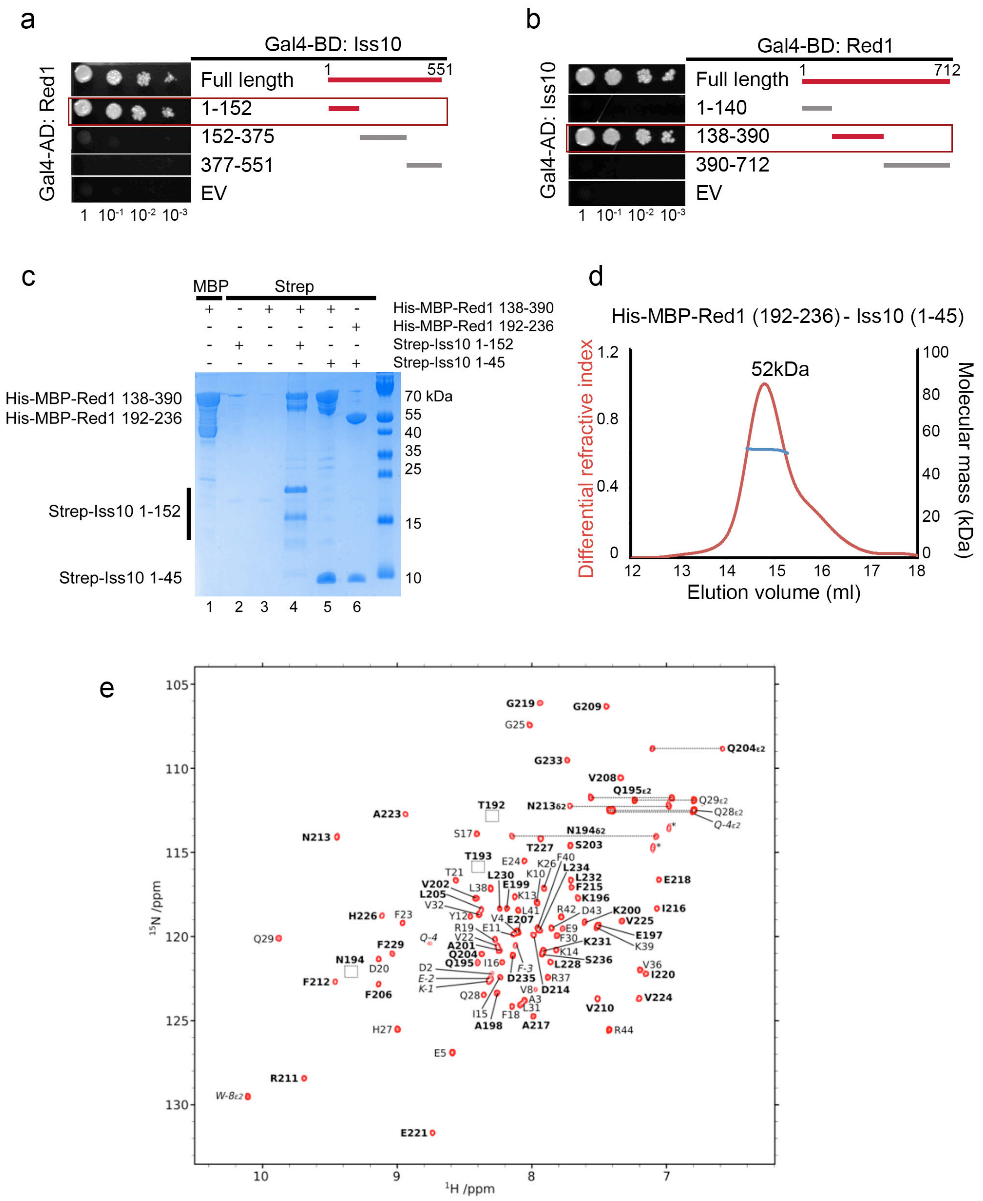

**Supplementary Figure 1**

**(a)** Mapping of the Iss10 interacting regions with Red1 by Y2H assays, revealing the Iss10 N-terminus to be sufficient for the interaction. Growth was assayed on medium lacking histidine and with 50mM 3AT. EV – empty vector**.** **(b)** Mapping of the Red1 interacting regions with Iss10 by Y2H assays, showing that the Red1 middle segment mediates the interaction. **(c)** SDS-PAGE analysis of Strep-tag pull-down experiments with Red1 and Iss10. The indicated combinations of His-MBP-Red1 and Strep-tagged Iss10 constructs were co-expressed and then purified on Strep-Tactin columns. His-MBP-Red1$^{138\text{-}390}$ purified alone on Amylose resin is shown as a control in lane 1. **(d)** Molecular mass determination of the His-MBP-Red1$^{192\text{-}236}$ and Strep-Iss10$^{1\text{-}45}$ complex by MALLS. The measured molecular masses of 52 kDa corresponds to a 1:1 stoichiometry. Calculated Mw of the complex is 56 kDa (49.4 + 6.7 kDa). The complex was injected at 2 mg/ml. **(e)** Annotated $^{15}$N-HSQC spectrum of the Iss10-Red1 heterodimer. The spectrum was collected at 298 K at a field strength of 800 MHz on a sample of 150 µM [$^{15}$N]Iss10-Red1 in 20 mM Tris (pH 7.5), 150 mM NaCl, 5 mM beta-mercaptoethanol and 10% (v/v) $D_2O$. Amide $^{1}$H,$^{15}$N crosspeaks are annotated by residue type and number. Residues in normal type are from Iss10, and in bold type for Red1. Annotations in italics are for residues in the expression tag, and unassigned side chain amides are indicated with an asterisk. Squares indicate the location of amide crosspeaks with low intensity due to line-broadening.

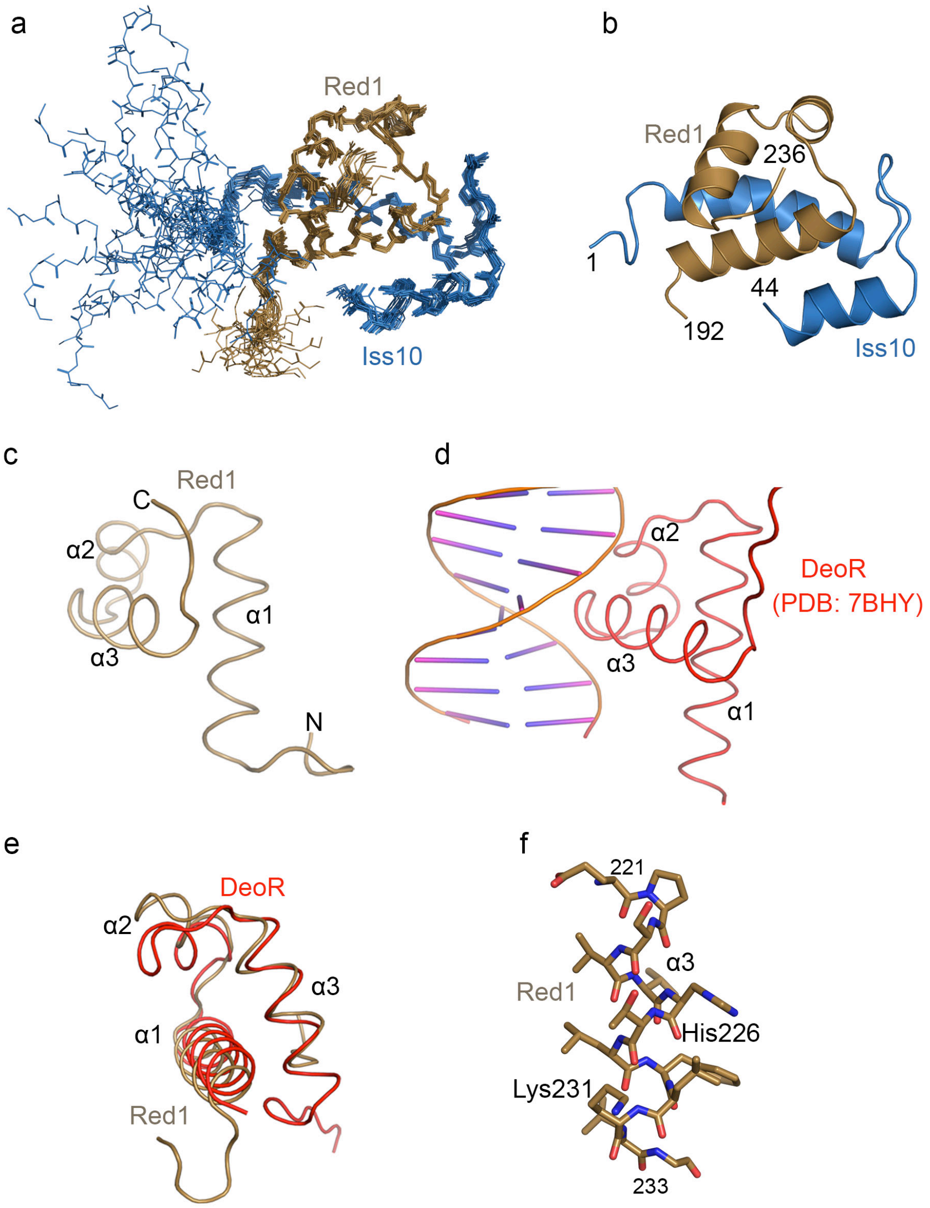

**Supplementary Figure 2**
**(a)** Superposition of 20 energy-minimized conformers representing the 3D NMR structure of the complex between Red1$^{192\text{-}236}$ and Iss10$^{1\text{-}45}$. Red1 is shown in brown and Iss10 in blue. **(b)** Cartoon representation of the structured part of the complex including Red1 residues 192-236 and Iss10 residues 1-44. **(c)** The HTH domain of Red1. **(d)** The HTH domain of *B. subtilis* DeoR in complex with dsDNA[1]. The structure was superimposed on Red1 shown in **c** with r.m.s.d. of 2.3 Å for 74 Cα atoms. **(e)** Superposition of HTH motifs of Red1 and DeoR. **(f)** Detail view of side chains of the Red1 helix α3.

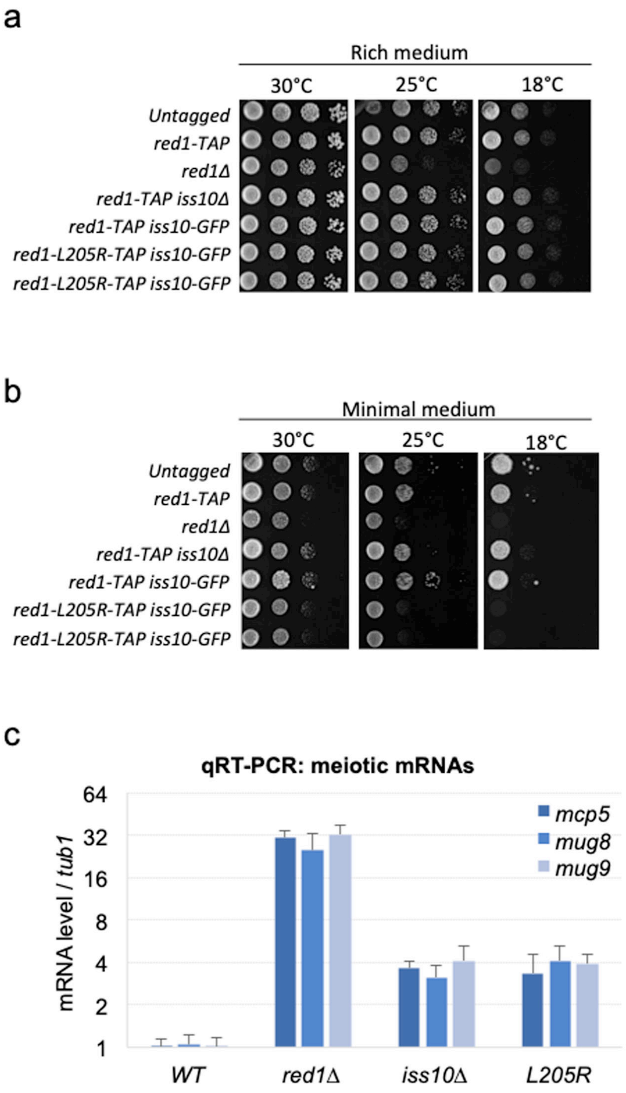

**Supplementary Figure 3**
**(a,b)** Growth assays comparing the proliferation of *red1-L205R-TAP* mutant cells to the parental cells, expressing the endogenous and untagged Red1 (*Untagged*), to cells expressing the wild-type Red1 fused to the TAP tag (*red1-TAP* or *WT*), to cells deleted for *red1* coding sequence (*red1*Δ), and to cells deleted for *iss10* coding sequence (*iss10*Δ). Cells were grown on rich (**a**) or minimal (**b**) medium, at temperatures of 30, 25 or 18ºC. **(c)** Quantitative RT-PCR showing the level of *mcp5*, *mug8* and *mug9* mRNAs normalized to *tub1* mRNA in *WT* and *red1-L205R* mutant cells. Total RNAs were purified from exponentially growing cells in liquid rich medium.

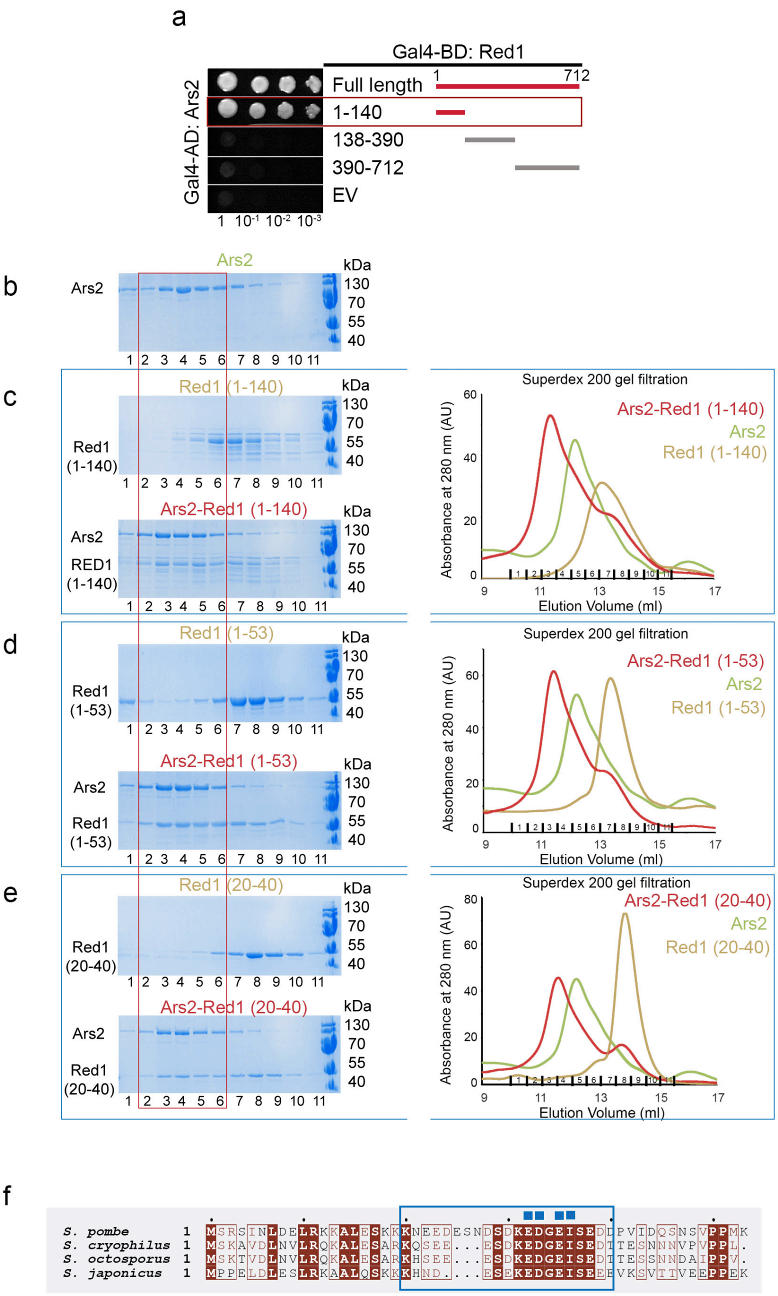

**Supplementary Figure 4**

**(a)** Y2H assays revealed that the N-terminal part of Red1 mediates the interaction with Ars2. Growth was assayed on medium lacking histidine and with 20mM 3AT. EV – empty vector. **(b)** SDS-PAGE analysis of fractions 1-11 of the Superdex 200 gel filtration elution profile of FL His-MBP-Ars2.

**(c)** Left panel - SDS-PAGE analysis of fractions 1-11 of the S200 gel filtration elution profiles of His-MBP-Red1$^{1\text{-}140}$ and its complex with FL His-MBP-Ars2. Right panel: overlay of Superdex 200 gel filtration elution profiles of FL His-MBP-Ars2, His-MBP-Red1$^{1\text{-}140}$ and their complex. The proteins were expressed individually as MBP fusions and purified on amylose resin. Both proteins were first purified on Superdex 200. **(d)** Superdex 200 gel filtration profiles and SDS-PAGE analysis of eluted fractions as described above, of His-MBP-Red1$^{1\text{-}53}$ and its complex with FL His-MBP-Ars2. **(e)** Superdex 200 gel filtration profiles and SDS-PAGE analysis of eluted fractions as described above, of His-MBP-Red1$^{20\text{-}40}$ and its complex with FL His-MBP-Ars2. **(f)** Sequence alignment of the Red1 proteins from different *Schizosaccharomyces* species. Only the sequence of residues 1-53 covering the N-terminal fragment involved in the interaction with Ars2 is shown. The minimal binding segment identified above is highlighted with a blue rectangle. Identical residues are in brown boxes. Blue squares indicate residues whose mutation impacts the interaction as shown in Fig. 5d-f.

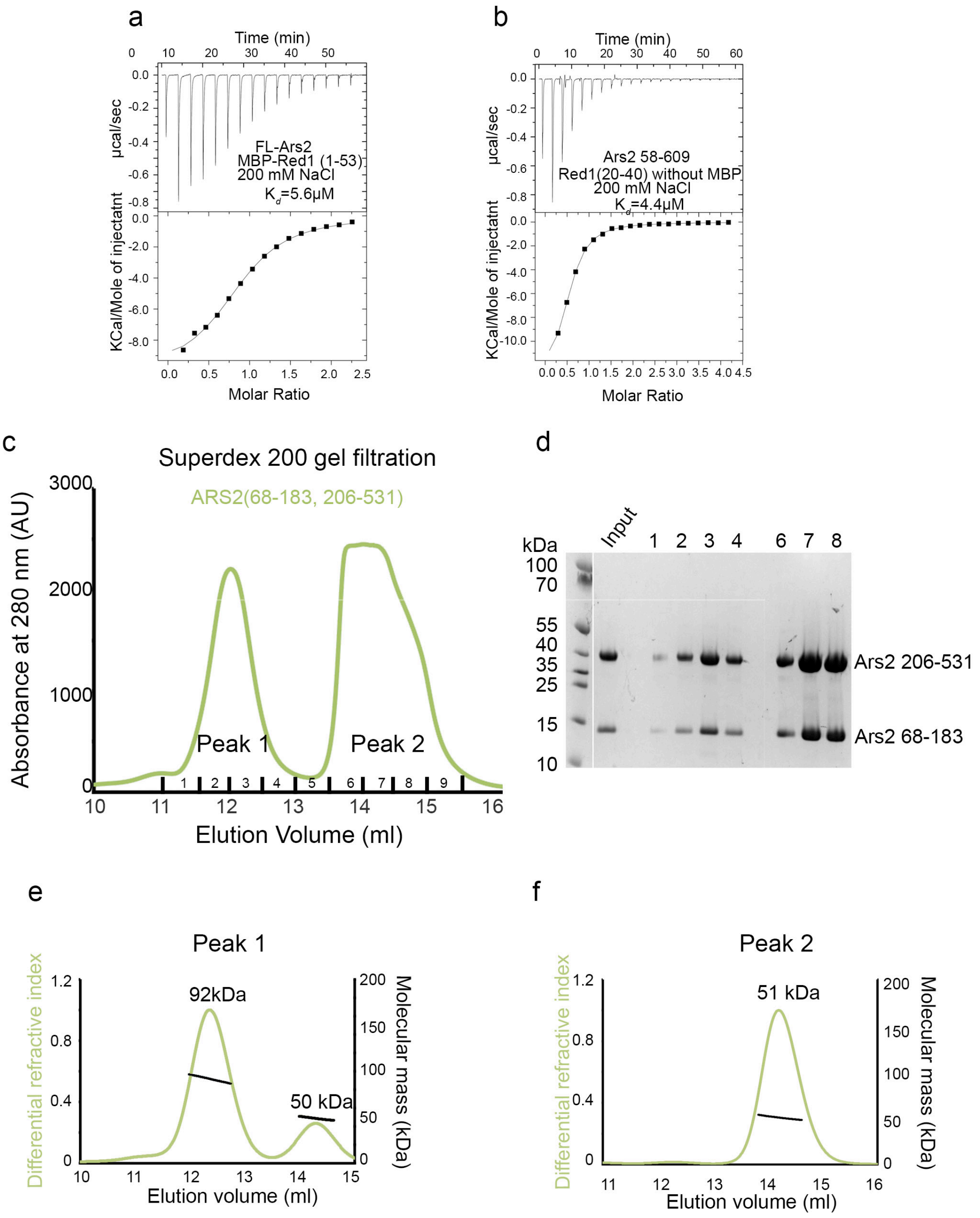

**Supplementary Figure 5**

**(a)** ITC measurement of the interaction affinity between FL Ars2 and His-MBP-Red1$^{1\text{-}53}$ in presence of 200mM NaCl. **(b)** ITC measurement of the interaction affinity between Ars2$^{58\text{-}609}$ and Red1$^{20\text{-}40}$ peptide in presence of 200mM NaCl. **(c)** Superdex 200 gel filtration elution profile of Ars2$^{68\text{-}183,206\text{-}531}$ revealing two elution peaks. **(d)** SDS-PAGE analysis of fractions 1-11 of the Superdex 200 gel filtration elution profile of Ars2$^{68\text{-}183,206\text{-}531}$ shown in **a**. **(e)** Molecular mass determination of the *S. pombe* Ars2$^{68\text{-}183,206\text{-}531}$ originating from the first elution peak shown in **a** by MALLS. The measured molecular masses of 92 and 50 kDa correspond to its dimer and monomer, respectively. The protein was injected at 38 mg/ml. **(f)** MALLS measurement of the molecular mass of the *S. pombe* Ars2$^{68\text{-}183,206\text{-}531}$ eluting in the second peak shown in **a**. The measured molecular mass of 51 kDa corresponds to a monomer. The protein was injected at 4 mg/ml.

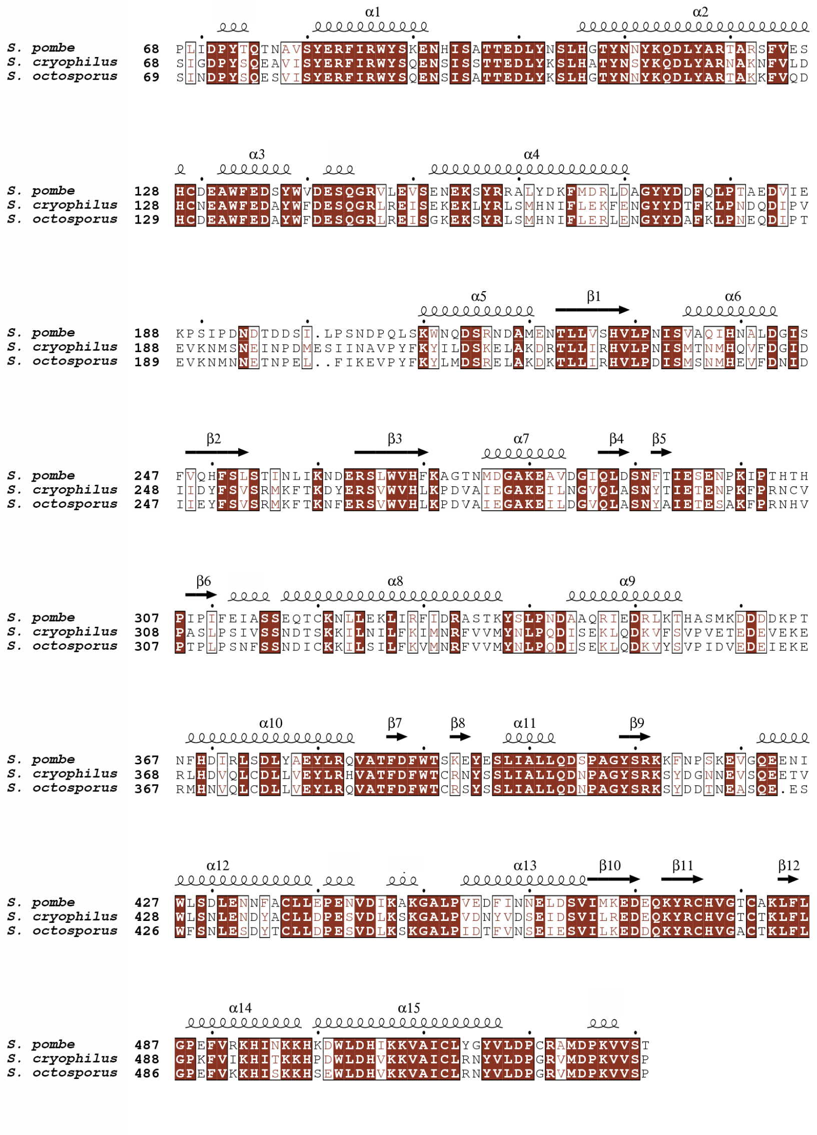

**Supplementary Figure 6**

Sequence alignment of Ars2 proteins from *Schizosaccharomyces* species. Identical residues are in brown boxes.

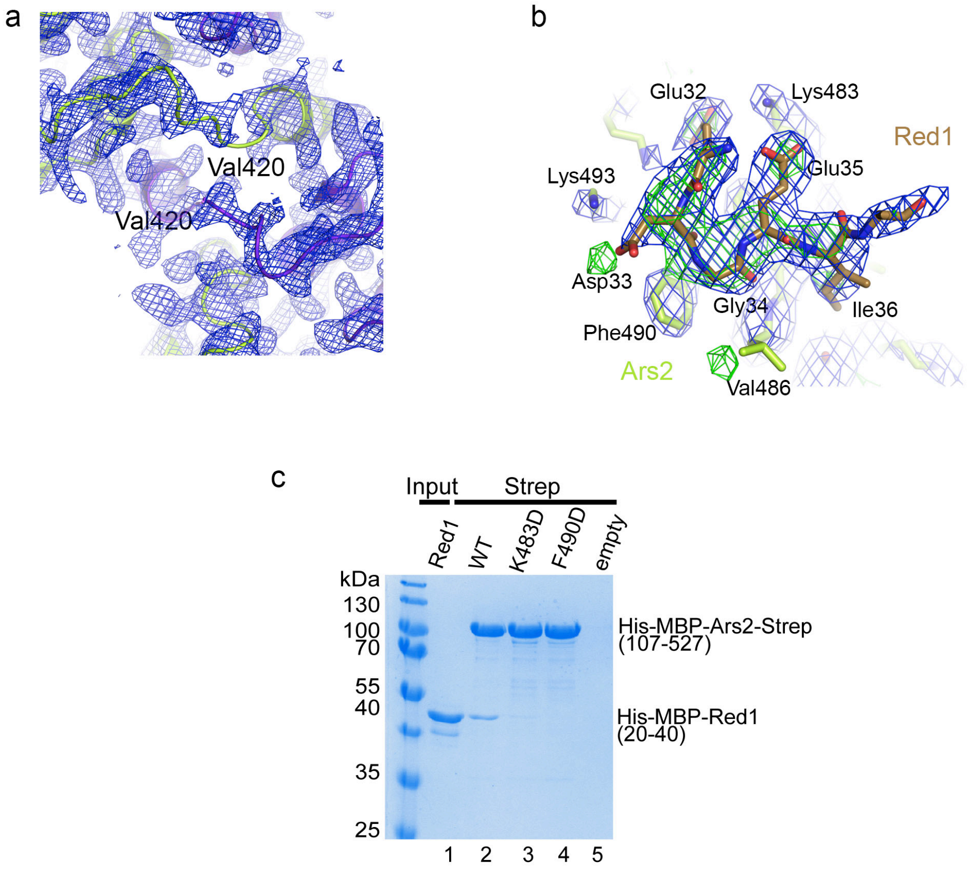

**Supplementary Figure 7**

**(a)** A 2*F*o-*F*c electron density map covering the two hinge regions around Val420 mediating the swap of the C-terminal domain between the two Ars2 protomers contoured at 1.0 σ. **(b)** 2*F*o-*F*c (blue) and omit *F*o-*F*c (green) electron density maps covering the Red1 residues interacting with Ars2 contoured at 1.0 and 2.5 σ, respectively. **(c)** Strep-tag pull-down experiments with His-MBP-$Red1^{20\text{-}40}$ and Strep-$Ars2^{107\text{-}527}$ mutants indicated above the lanes. Ars2 and Red1 proteins were first purified using $Ni^{2+}$ resin (lane 1). Red1 with Ars2 variants were then co-purified on Strep-tactin resin (lanes 2-4).

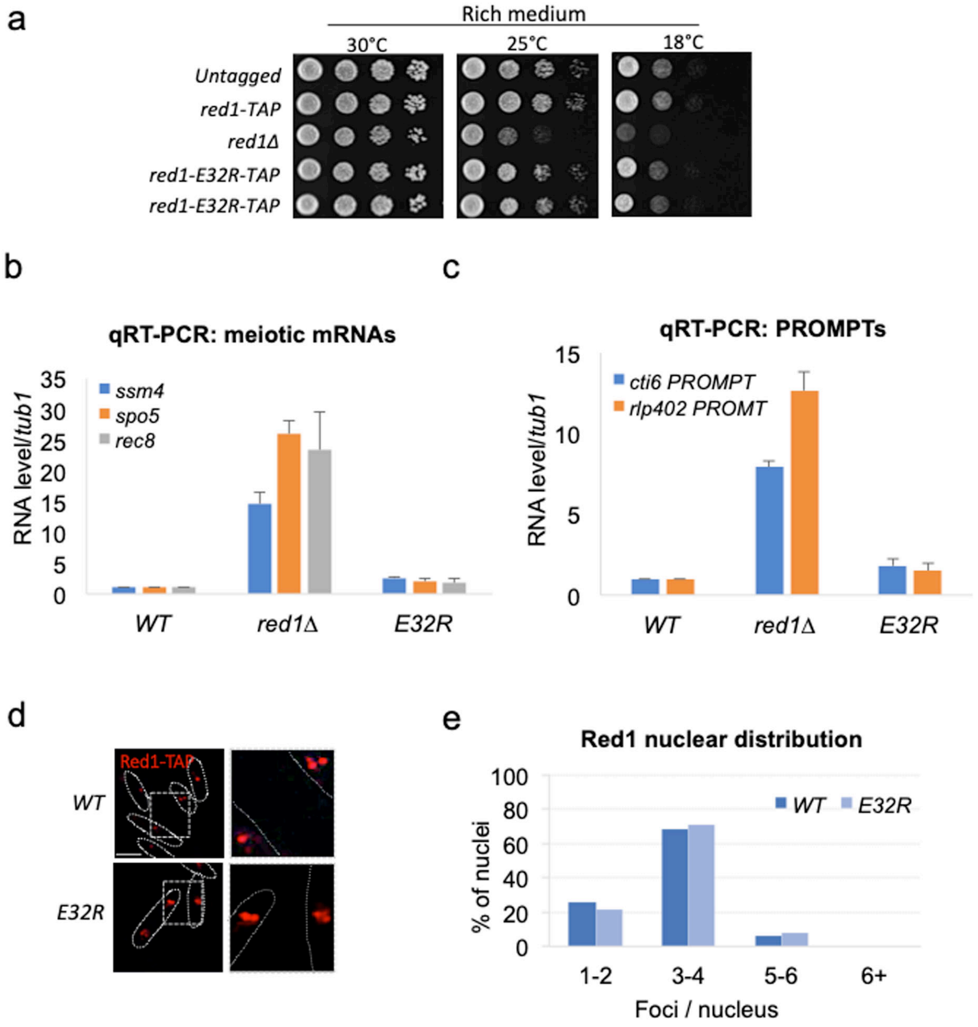

**Supplementary Figure 8**

**(a)** Growth assays comparing the growth of *red1-E32R-TAP* mutant cells to the parental cells, which express endogenous and untagged Red1 (*Untagged*), to cells expressing wild-type Red1 fused to the TAP tag (*red1-TAP* or *WT*), to cells deleted for *red1* coding sequence (*red1*Δ). Cells were grown on rich medium, at temperatures of 30, 25 or 18ºC. **(b)** Quantitative RT-PCR monitoring the level of meiotic DSR-containing mRNAs (*ssm4*, *spo5*, *rec8*) and **(c)** of PROMPTs-CUTs (*cti6* and *rlp402* PROMPTs) in *WT* and *red1-L205R* mutants. Total RNAs were purified from exponentially growing cells in liquid minimal medium. Variation of RNA levels is expressed in fold-change relative to the *WT* levels and normalized to *tub1* mRNA. **(d)** Images of immuno-fluorescence microscopy showing Red1-TAP localization in *WT* and *red1-E32R* cells. Scale bar = 5 µm. **(e)** Graph representing the foci number per nucleus of Red1-TAP in *WT* and *red1-E32R* cells. Count performed on 100 nuclei from two different biological isolates.

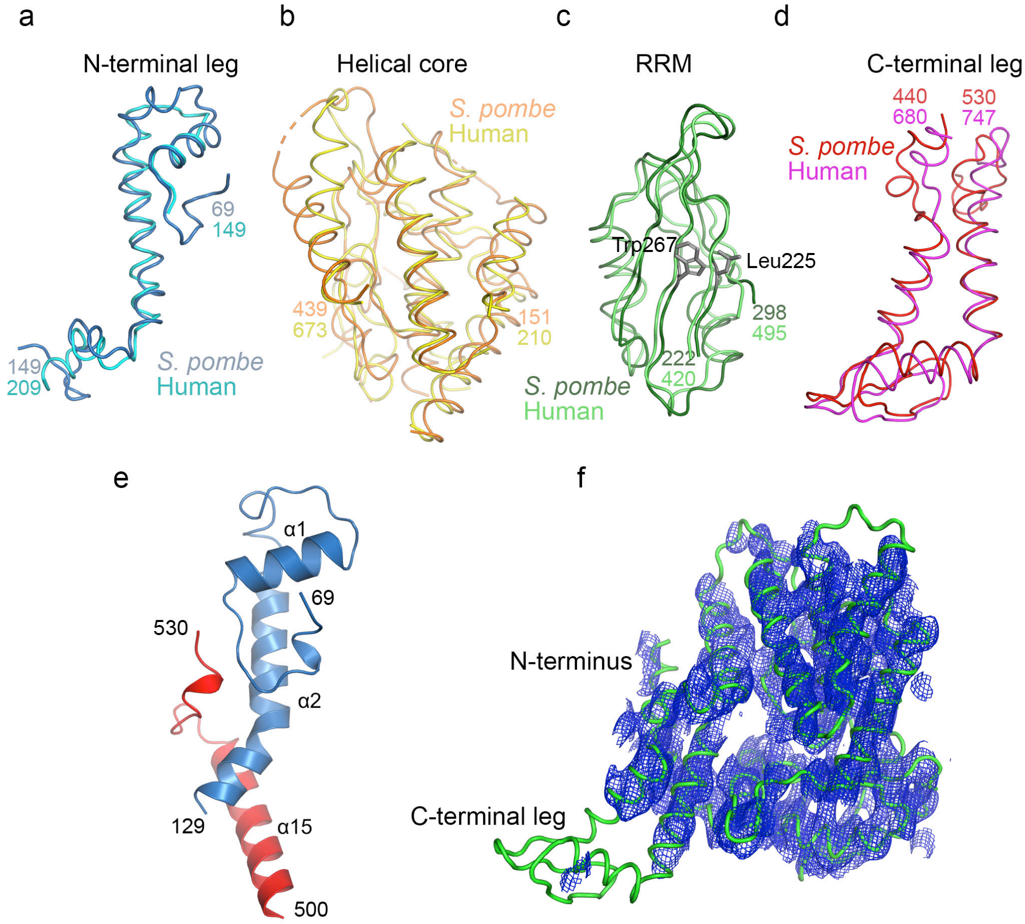

**Supplementary Figure 9**

**(a)** Comparison of *S. pombe* and human Ars2 N-terminal leg (r.ms.d.- 2.2 Å for 52 Cα atoms).
**(b)** Comparison of *S. pombe* and human Ars2 helical core (r.ms.d.- 2.8 Å for 153 Cα atoms).
**(c)** Comparison of *S. pombe* and human Ars2 RRM domain (r.ms.d.- 1.6 Å for 74 Cα atoms).
**(d)** Comparison of *S. pombe* and human Ars2 C-terminal leg (r.ms.d.- 4.2 Å for 60 Cα atoms). **(e)** The N-terminal segment (residues 69-80) packs against helices α1 and α2 and interact with C-terminal residues 527-530. **(f)** A 5 Å resolution 2*F*o-*F*c electron density map covering a monomer of Ars2$^{107-527}$. The N-terminal helix and Zn-finger domain of the C-terminal leg are disordered. The structure was determined by molecular replacement using the Alphafold *S. pombe* Ars2 model (AF-O94326). The model bias was reduced using the Prime-and-switch phasing in Resolve[2].

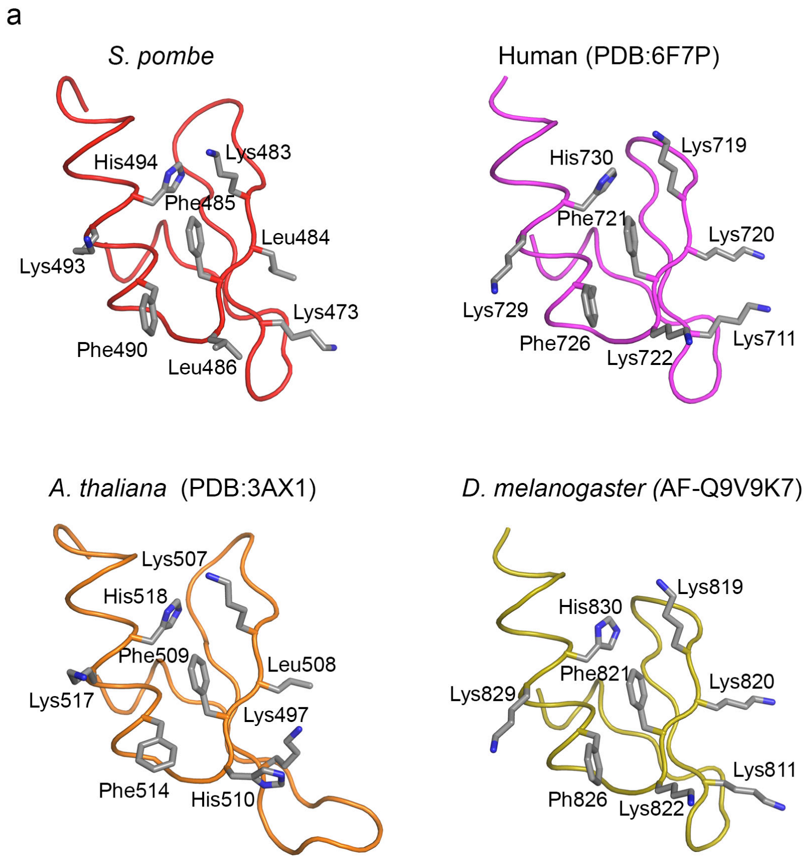

**Supplementary Figure 10**

**(a)** Comparison of the conserved Red1-binding surface among *S. pombe* (this study), human (PDB code-6F7P) and *A. thaliana* (PDB code-3AX1) structures and the Aplhafold model of *D. melanogaster* ARS2 (AF-Q9V9K7).

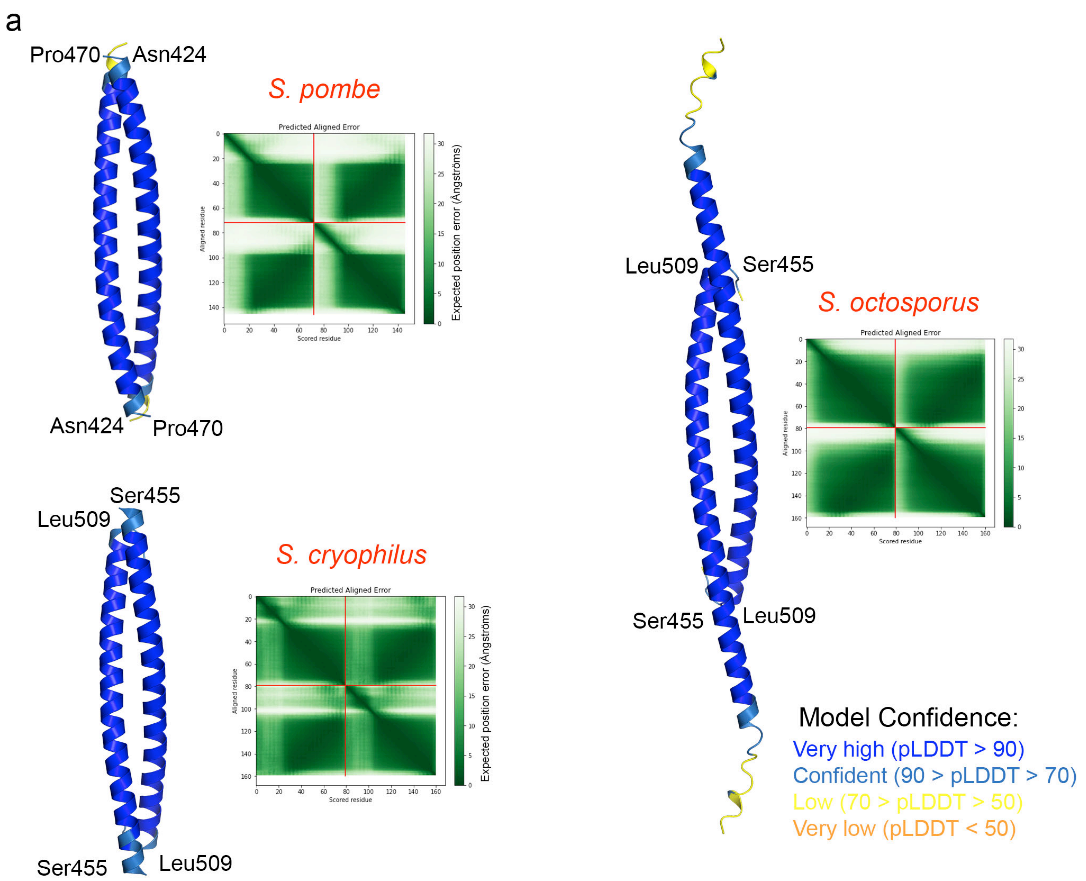

**Supplementary Figure 11**

**(a)** Alphafold models of the *S. pombe, S. octosporus and S. cryophilus* Red1 C terminus. Only the dimeric coiled-coil regions, corresponding to the alignment in Fig. 7d are shown. For *S. octosporus*, longer chains, predicted with high confidence are shown. The structures are colored according to the Aplhafold per-residue confidence score (pLDDT). Predicted aligned error plot is shown for each model.

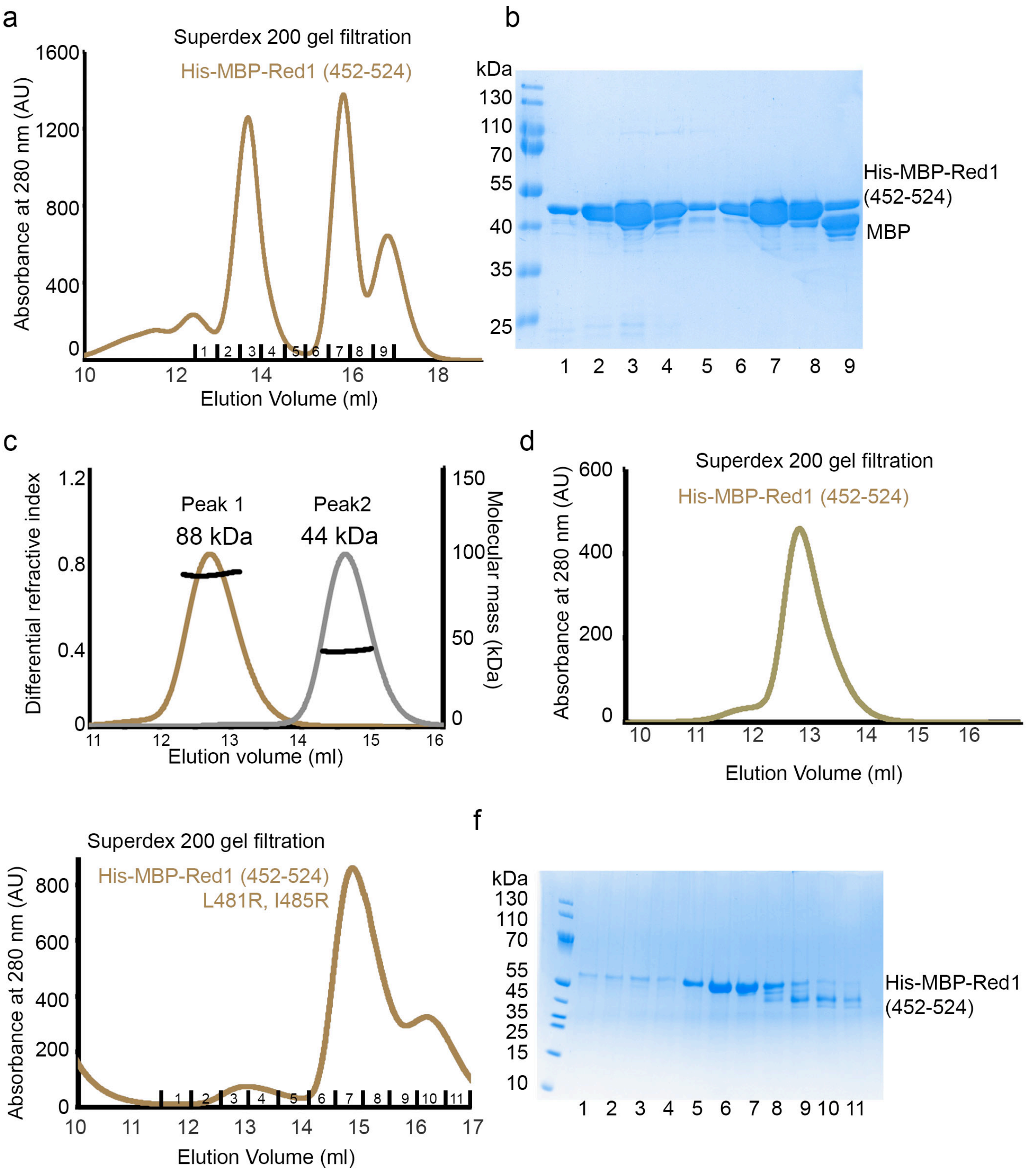

**Supplementary Figure 12**

**(a)** Superdex 200 gel filtration elution profile of His-MBP-Red1$^{452-524}$ revealing two elution peaks (third peak corresponds to MBP). **(b)** SDS-PAGE analysis of fractions 1-10 of the Superdex 200 gel filtration elution profile of His-MBP-Red1$^{452-524}$ shown in **a**. **(c)** Molecular mass determination of the His-MBP-Red$^{452-524}$ by MALLS. The measured molecular mass of 88 kDa and 44 kDa correspond to a dimer and monomer of Red1 (peak 1 and 2 of gel filtration shown in **a**). Calculated molecular mass of a monomer is 53 kDa. The two samples were injected at 9 and 5 mg/ml, respectively. **(d)** 2$^{nd}$ round Superdex 200 gel filtration of His-MBP-Red1$^{452-524}$ originating from the first elution peak shown in **a.** **(e)** Superdex 200 gel filtration profile of His-MBP-Red1$^{452-524}$ (L481R, I485R) mutant. The protein mostly elutes in a peak corresponding to the monomer. A large portion of the mutated protein also elutes in the void volume, indicating that monomeric His-MBP-Red1$^{452-524}$ becomes unstable. **(f)** SDS-PAGE analysis of fractions 1-11 of the Superdex 200 gel filtration elution profile of His-MBP-Red1$^{452-524}$ (L481R, I485R) shown in **e**.

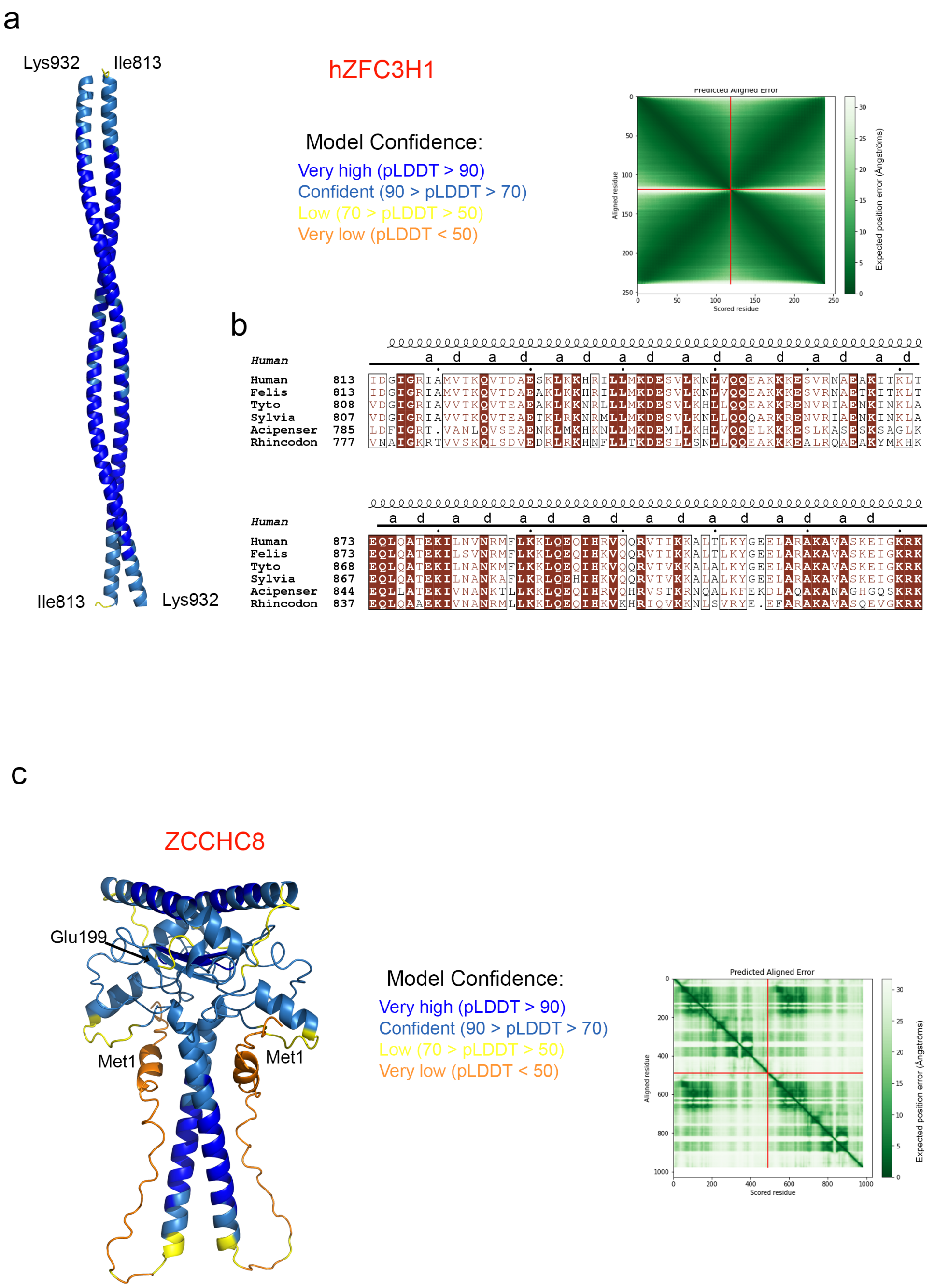

**Supplementary Figure 13**

**(a)** Alphafold model of human ZFC3H1 dimer. Only the dimeric coiled-coil regions, corresponding to the alignment shown in **b** are shown. The structure is colored according to the Alphafold pLDDT score. Predicted aligned error plot is shown. **(b)** Sequence alignment of ZFC3H1 proteins. Only the sequence of the coiled-coil region is shown. Identical residues are in brown boxes, and conserved residues are shown in brown. Heptad positions are indicated. **(c)** Alphafold model of human ZCCHC8 dimer. Only residues 1-199 are shown, that were predicted with confidence of mutual positions of the two protomers according to the predicted aligned error plot. The structure is colored according to the Alphafold pLDDT score.

**Supplementary Table 1. NMR and refinement statistics for the minimal heterodimer Iss10/Red1.**

| | Iss10/Red1 |
|---|---|
| **Number of structures in ensemble** | 20 |
| **Distance Constraints** | |
| Total | 1993 |
| Protein Intra-residue | 692 |
| Protein Inter-residue | |
| Sequential (\|i-j\|=1) | 321 |
| Short Range (1<\|i-j\|<4) | 142 |
| Medium Range (3<\|i-j\|<6) | 79 |
| Long Range (\|i-j\|>5) | 74 |
| Intermolecular | 87 |
| Ambiguous | 528 |
| Hydrogen bond | 70 |
| **Total dihedral angle constraints** | |
| Backbone ($\varphi$,$\psi$) | 154 (77,77) |
| Sidechain ($\chi_1$,$\chi_2$) | 58 (46,12) |
| **Structure Statistics** | |
| **Violations (mean and SD)** | |
| Distance Constraints (Å)[a] | 0.023 ± 0.002 |
| Dihedral Angle Constraints (°)[b] | 2.28 ± 0.09 |
| RDC, Q[c] | 0.24 ± 0.01 |
| **Deviations from idealized geometry** | |
| Bond Lengths (Å) | 0.003 ± 0.000 |
| Bond Angles (°) | 0.45 ± 0.01 |
| Impropers (°) | 1.2 ± 0.1 |
| **Ramachandran plot (%)[d]** | |
| Most favored | 92.9 (88.2) |
| Additionally favored | 6.6 (10.7) |
| Generally allowed | 0.4 (0.7) |
| Disallowed | 0.1 (0.4) |
| **Average pairwise rmsd (Å)** | |
| Protein backbone 2° structure | 0.5 ± 0.2 |
| Protein heavy 2° structure | 1.0 ± 0.2 |
| Protein backbone all | 1.2 ± 0.3 |
| Protein heavy all | 1.3 ± 0.2 |

[a] No distance violation greater than 0.5 Å.
[a] No dihedral restraint violation greater than 10°.
[c] Calculated for each structure in the ensemble using the method described by [3].
[d] Determined by using PROCHECK-NMR[4]. Values do not include the expression tag. Values that include the tag are in parentheses.

**Supplementary Table 2. Crystallographic data collection and refinement statistics.**

| | **Red1-Ars2** |
|---|---|
| **Data collection** | |
| Space group | $P2_1$ |
| Cell dimensions | |
| *a*, *b*, *c* (Å) | 64.7, 128.3, 89.3 |
| α β γ (°) | 90, 108, 90 |
| Resolution (Å) | 85-2.8 (3-2.8)[a] |
| $R_{merge}$ (%) | 29.6 (148) |
| I / σI | 5.3 (1.5) |
| $CC_{½}$ (%) | 99 (55) |
| Completeness (%) | 76 (18) |
| Redundancy | 5.6 (6.2) |
| **Refinement** | |
| Resolution (Å) | 84.8-2.8 |
| No. reflections | 25473 |
| $R_{work}$ / $R_{free}$ | 0.245/0.302 |
| *B*-factors | 53 |
| R.m.s. deviations | |
| Bond lengths (Å) | 0.002 |
| Bond angles (°) | 1.17 |

[a] Values in parentheses are for highest resolution shell.

### Supplementary Table 3. List of *S. pombe* strains used in this study.

| Name | Genotype | Figures | Source |
|---|---|---|---|
| SPV8 | *h+ leu1-32 ori1 ade6 M216 ura4-D18 imr1R::ura4+* | Sup3, sup8 | Lab Stock |
| SPV2180 | *h+ leu1-32 ori1 ade6 M216 ura4-D18 imr1R::ura4+ red1::TAP KanR* | Fig3,5 & sup3, 8 | Lab Stock |
| SPV3106 | *h+ leu1-32 ori1 ade6 M216 ura4-D18 imr1R::ura4+ red1Δ::hphR* | Fig3,5 & sup3, 8 | Lab Stock |
| SPV5699 | *h+ leu1-32 ori1 ade6 M216 ura4-D18 red1::TAP KanR* | Sup3, sup8 | This study |
| SPV5714 | *h+ leu1-32 ori1 ade6 M216 ura4-D18 red1Δ30-207::TAP KanR* | | This study |
| SPV5785 | *h+ leu1-32 ori1 ade6 M216 ura4-D18 red1E32R::TAP KanR Clone 29* | Fig5, sup8 | This study |
| SPV5786 | *h+ leu1-32 ori1 ade6 M216 ura4-D18 red1E32R::TAP KanR Clone 30* | Fig5, sup8 | This study |
| SPV5787 | *h+ leu1-32 ori1 ade6 M216 ura4-D18 red1L205R::TAP KanR Clone 4* | Fig3, sup3 | This study |
| SPV5788 | *h+ leu1-32 ori1 ade6 M216 ura4-D18 red1L205R::TAP KanR Clone 14* | Fig3, sup3 | This study |
| SPV5994 | *h- leu1-32 ori1 ade6 M216 ura4-D18 imr1R::ura4+ iss10Δ::hphR* | Sup3 | This study |
| SPV5995 | *h+ leu1-32 ori1 ade6 M216 ura4-D18 imr1R::ura4+ iss10Δ::hphR* | Sup3 | This study |
| SPV6013 | *h- leu1-32 ori1 ade6 M216 ura4-D18 imr1R::ura4+ red1L205R::TAP KanR iss10Δ::hphR Clone 3* | Fig3, sup3 | This study |
| SPV6015 | *h+ leu1-32 ori1 ade6 M216 ura4-D18 imr1R::ura4+ red1L205R::TAP KanR iss10Δ::hphR Clone 7* | Fig3, sup3 | This study |
| SPV6017 | *h+ leu1-32 ori1 ade6 M216 ura4-D18 red1::TAP KanR iss10::GFP NatR Clone 5* | Fig3, sup3 | This study |
| SPV6018 | *h+ leu1-32 ori1 ade6 M216 ura4-D18 red1::TAP KanR iss10::GFP NatR Clone 6* | Fig3, sup3 | This study |
| SPV6020 | *h+ leu1-32 ori1 ade6 M216 ura4-D18 red1L205R::TAP KanR iss10::GFP NatR Clone 5* | Fig3, sup3 | This study |
| SPV6021 | *h+ leu1-32 ori1 ade6 M216 ura4-D18 red1L205R::TAP KanR iss10::GFP NatR Clone 6* | Fig3, sup3 | This study |
| SPV6023 | *h- leu1-32 ori1 ade6 M216 ura4-D18 imr1R::ura4+ ars2-GFP Clone 1* | sup8 | This study |
| SPV6024 | *h- leu1-32 ori1 ade6 M216 ura4-D18 imr1R::ura4+ ars2-GFP Clone 2* | sup8 | This study |
| SPV6026 | *h+ leu1-32 ori1 ade6 M216 ura4-D18 red1E32R::TAP KanR ars2-GFP Clone 1* | Fig5, sup8 | This study |
| SPV6027 | *h+ leu1-32 ori1 ade6 M216 ura4-D18 red1E32R::TAP KanR ars2-GFP Clone 2* | Fig5, sup8 | This study |
| SPV6029 | *h+ leu1-32 ori1 ade6 M216 ura4-D18 imr1R::ura4+ red1::TAP KanR ars2-GFP Clone 1* | Fig5, sup8 | This study |
| SPV6030 | *h+ leu1-32 ori1 ade6 M216 ura4-D18 imr1R::ura4+ red1::TAP KanR ars2-GFP Clone 2* | Fig5, sup8 | This study |

**Supplementary Table 4. List of primers used for the *S. pombe* analysis.**

| gene | Sequence 5' > 3' |
|---|---|
| **strain construction** | |
| *Ars2-GFP* tag for | TTCTATAGAACTTATCAAGATTTGGATGCGCCTAACCAGGAGGTTCCAGAATTAGATTACcggatccccgggttaattaa |
| *Ars2-GFP* tag rev | TAACGTCTATACAGGAAATGATATTAACCAAAAATCTTTCTGACTAAGAACATAAATTTAgaattcgagctcgtttaaac |
| *Iss10-GFP* tag for | AACGACTATGAATTAGCAGAGGAAATTGAGAGGCGTCGTCATCATAGCTTTTATGATTATcggatccccgggttaattaa |
| *Iss10-GFP* tag rev | TCAAATATTCCGCCAAGCAATTTTTAAATTGTTGCGCGCACTTACGGAAGAGTATAGCTAgaattcgagctcgtttaaac |
| *red1Δ30-207* for | GGTCAATAAATCTCGATGAGTTGCGAAAAAAGGCTTTAGAATCAAAAAAGAAGAATGAAGAAGATGAGAGTAATGACTCTCGCCAGGGTTTTCC CAGTCACGAC |
| *red1Δ30-207* rev | AGTCCGAGCTTTAAAAATAAAGTATGAACAACCGAAGGCTCGATGCCCTCAGCAATAAAATCGTTAAAGCGCACTCCTACAGCGGATAACAATT TCACACAGGA |
| **genotyping** | |
| *red1Δ30-207* for | TTGCTACATTCAGTATACCTGTTCA |
| *red1Δ30-207* rev | TTCGCTAATTTCGCCATCTT |
| *Ars2-GFP* tag for | GGATCCCAAAGTGGTAAGCA |
| *Ars2-GFP* tag rev | TTCTATCTGTGCAGCGTTCG |
| *Iss10-GFP* tag for | GCCTCCTATTGAAACGCACT |
| *Iss10-GFP* tag rev | TGCGAAACCAATTTATGCAA |
| **qPCR** | |
| *tub1* for | GTACTGGCCCATACCGTGAT |
| *tub1* rev | CGAATGGAAGACGAGAAAGC |
| *ssm4* for | TGCAAGAGGAAACTCAAAGG |
| *ssm4* rev | TTCCTCCTCCACTTGTTTTGA |
| *spo5* for | TTATGGCGTGCTCGTGAATA |
| *spo5* rev | TTCGACTCCAATGGCACATA |
| *mug8* for | GGCTGACCAAATGAATGCTT |
| *mug8* rev | GATCTGCCCAACCAACAATC |
| *mug9* for | CTCTGTTGGCTCCTTCAAGC |
| *mug9* rev | TAATGTTTGAGCACGCATGG |
| *mcp5* for | TTTGGATTAACCCGCATTGT |
| *mcp5* rev | TGGCGTTCTTTAAGGCATCT |
| *rec8* for | GTTGAAGTTGGACGGGATGT |
| *rec8* rev | TTCTACCCTACTCGGCATCG |
| *cti6 PROMPT* for | CATCGGAGAACCTGGTTTGT |
| *cti6 PROMPT* rev | TGCATTCCCAAATCAAGTC |
| *rlp402 PROMPT* for | GAAAGAGGAACAATGTTTAGAAGTC |
| *rlp402 PROMPT* rev | AAGCCCTACGATTGCTGTG |
